# A Bayesian analysis of birth pulse effects on the probability of detecting Ebola virus in fruit bats

**DOI:** 10.1101/2023.08.10.552777

**Authors:** David R.J. Pleydell, Innocent Ndong Bass, Flaubert Auguste Mba Djondzo, Dowbiss Meta Djomsi, Charles Kouanfack, Martine Peeters, Julien Cappelle

## Abstract

Since 1976 various species of Ebolavirus have caused a series of zoonotic outbreaks and public health crises in Africa. Bats have long been hypothesised to function as important hosts for ebolavirus maintenance, however the transmission ecology for these viruses remains poorly understood. Several studies have demonstrated rapid seroconversion for ebolavirus antibodies in young bats, yet paradoxically few PCR studies have confirmed the identity of the circulating viral species causing these seroconversions. The current study presents an age-structured epidemiological model that characterises the effects of seasonal birth pulses on ebolavirus transmission within a colony of African straw-coloured fruit bats (*Eidolon helvum*). Bayesian calibration is performed using previously published serological data collected from Cameroon, and age-structure data from Ghana. The model predicts that annual birth pulses most likely give rise to annual outbreaks, although more complex dynamic patterns – including skip years, multi-annual cycles and chaos – may be possible. Weeks 30 to 31 of each year were estimated to be the most likely period for isolating the circulating virus in Cameroon. The probability that a previous PCR campaign failed to detect Ebola virus, assuming that it was circulating, was estimated to be one in two thousand. This raises questions such as (1) what can we actually learn from ebolavirus serology tests performed without positive controls? (2) are current PCR tests sufficiently sensitive? (3) are swab samples really appropriate for ebolavirus detection? The current results provide important insights for the design of future field studies aiming to detect Ebola viruses from sylvatic hosts, and can contribute to risk assessments concerning the timing of zoonotic outbreaks.

## Introduction

Bats have been implicated as reservoir hosts to numerous viruses of zoonotic or animal health importance, including: Hendra virus, Marburg virus, Middle East respiratory coronavirus (MERS-CoV), Nipah virus, severe acute respiratory syndrome coronavirus (SARS-CoV), SARS-CoV-2 and Swine acute diarrhoea syndrome corona-virus (Letko et al., 2020). This list of bat-borne emerging viruses is thought to also include filoviruses of the *Ebolavirus* genus (Caron et al., 2018; Feldmann et al., 2020; Leroy et al., 2005). Indeed, the filovirus-like VP35 gene is estimated to have been maintained in bat genomes for 13.4 million years (Taylor et al., 2011), which provides evolutionary support for the long-term exposure of bats to filoviruses. However, despite numerous outbreaks of Ebola in equatorial Africa since 1976 – where fatality rates typically fall in the range of 40-70% (Jacob et al., 2020; Munster et al., 2018) – the hypotheses that (i) bats provide sylvatic reservoirs for Ebola viruses, and (ii) that these reservoirs contribute to spillover events, remain unconfirmed. Moreover, the eco-epidemiology of Ebola virus remains poorly understood, and empirical evidence for bats functioning as primary maintenance reservoirs for Ebola viruses remains non-conclusive (Olival and Hayman, 2014).

Serological data shows that some bat species express high seroprevalence for Ebola virus (De Nys et al., 2018; Hayman, Yu, et al., 2012). But serology is hard to interpret in bats without positive controls, and there is a paradoxal discrepancy between serological data and viral detection (Caron et al., 2018). Indeed, no Ebola virus has ever been isolated from bats, and only a few individuals of three bat species have tested positive by polymerase chain reaction (PCR) for Ebola virus (Leroy et al., 2005) – a result that remains to be replicated despite extensive sampling. Recent longitudinal monitoring of a straw-colored fruit bat (*Eidolon helvum*) population in Cameroon has shown extensive seroconversion of young (juvenile and sexually immature adult) bats over a period of a few months, suggesting active Ebola virus circulation - however, no bat tested positive for Ebola virus by PCR during that study (Djomsi et al., 2022). Another *E. helvum* study in Guinea provided similar results, with seroprevalence decreasing over the first months of life and increasing again in the first years of adult life, but again, no bats were found to be PCR positive (Champagne et al., Submitted).

Modelling is being increasingly used to help understand the interplay between ecological and epidemiological dynamics in bats (Glennon et al., 2019; Hayman, 2015; Peel, KS Baker, Hayman, Broder, et al., 2018). Considerable attention has been paid to the effects of seasonal birth pulses on the pool of susceptible individuals and subsequent epidemiological consequences (Hranac et al., 2019; Peel, Pulliam, et al., 2014). For example, strong seasonal patterns in the prevalence of rabies in bats have been attributed to epidemiological consequences of birth pulses (George et al., 2011), and the biannual birth pulses of some Egyptian fruit bat populations are thought to increase the probability of pathogen maintenance (Hayman, 2015). Modelling has also indicated that maternally-derived antibodies can contribute to viral maintenance (Hayman, Luis, et al., 2018).

In order to explore the enigmatic discrepancy between Ebola serology and virology data, we developed an age-structured epidemiological model that included seasonal birth pulses and waning immunity, and used Bayesian techniques to fit the model to longitudinal *E. helvum* serology data from Cameroon (Djomsi et al., 2022). Our three main objectives were as follows. First, to quantify uncertainty in the parameters and dynamics of the model given the seroprevalence data of Djomsi et al. (2022). Second, to quantify the probability of not detecting any PCR positive bats given the sampling scheme of the Djomsi et al. (2022) study. Third, to identify whether seasonal birthing patterns can help identify optimal time-windows for Ebola virus detection. This modelling work has identified potentially important biological parameters that can help explain the observed serology dynamics, and provides insights that can help improve the efficiency of surveillance strategies for detecting Ebola virus in bats. In particular, these analyses provide insights into practical questions concerning the establishment of adequate sampling efforts for virus isolation, and raise questions concerning the meaning of positive serological samples from bats.

## Material and methods

### Study site and eco-epidemiological data

Our analyses were based primarily on data from a longitudinal serology survey at an *E. helvum* colony in Yaounde, Cameroon (Djomsi et al., 2022). Although various antigens were used for serological testing in that study, we exclusively used data from the Res1GP.ZEBVkiss antigenic test – a test based on the glycoprotein of Zaire Ebola virus. We selected this data-set because, among all antigenic tests used, the Res1GP.ZEBVkiss test generated the highest seropositive rate and the strongest seasonal signal. This data provided information concerning seasonal variation in the presence of four different age classes: pups (*P*) - young non-weaned bats that remain attached to their mothers; juveniles (*J*) - weaned young, that do not yet display joint ossification; immature adults (*I*) - large bats with ossified joints but without any sign of sexual maturity; and adult bats (*A*). Pups were not sampled directly, however, lactating females provided a proxy for their presence. A summary of this data is provided in table 1.

**Table 1.** Summary of *E. helvum* serology and lactation data from Yaounde, Cameroon. Negative and positive results for the Res1GP.ZEBVkiss antigenic test are shown for captured bats of three age classes. The number of captured adult female bats either lactating or not lactating are also shown. Lactation was used as a proxy for inferring seasonality in the presence of pups. A full description of this data is available in Djomsi et al. (2022).

| Date | Juvenile |  | Immature |  | Adult |  | Female Adults |  |
| --- | --- | --- | --- | --- | --- | --- | --- | --- |
|  | Neg | Pos | Neg | Pos | Neg | Pos | Lactating | Not Lactating |
| 2018-12-07 | 0 | 0 | 2 | 0 | 10 | 8 | 0 | 8 |
| 2019-01-26 | 0 | 0 | 0 | 0 | 45 | 53 | 0 | 42 |
| 2019-03-03 | 0 | 0 | 0 | 0 | 10 | 7 | 1 | 7 |
| 2019-04-02 | 12 | 1 | 0 | 0 | 50 | 24 | 46 | 9 |
| 2019-05-07 | 116 | 1 | 0 | 0 | 21 | 10 | 20 | 6 |
| 2019-06-16 | 71 | 1 | 0 | 0 | 4 | 6 | 0 | 3 |
| 2019-07-17 | 13 | 4 | 67 | 10 | 27 | 11 | 0 | 16 |
| 2019-09-17 | 1 | 0 | 11 | 26 | 3 | 11 | 0 | 3 |
| 2019-10-15 | 0 | 0 | 16 | 22 | 15 | 16 | 0 | 11 |
| 2019-11-15 | 0 | 0 | 13 | 51 | 31 | 20 | 0 | 24 |

To help estimate adult mortality rates we used tooth cementum annuli data from 294 adult bats sampled in Ghana (Peel, KS Baker, Hayman, Suu-Ire, et al., 2016). Thus, we assume that the age structures at the sampled colonies in Ghana and Cameroon are equivalent.

### Mechanistic model

A system of ordinary differential equations was developed to provide a deterministic characterisation of Ebola transmission in an age-structured *E. helvum* population. This system is depicted graphically in figure 1 and algebraically in equations 1-19. A list of model parameters is presented in table 2. Age structure in the model was defined using the same four age classes recorded in the field (see above), namely: pups (*P*); juveniles (*J*); immature adults (*I*); and adult bats (*A*). Five epidemiological classes were used: protected by maternal antibodies (*M*); susceptible (*S*); infected (*I*); recovered (*R*); long-term immunity (*L*). For simplicity, it was assumed that each year is exactly 52 weeks long, and weeks are used as our time unit throughout (unless stated otherwise).

**Table 2.**
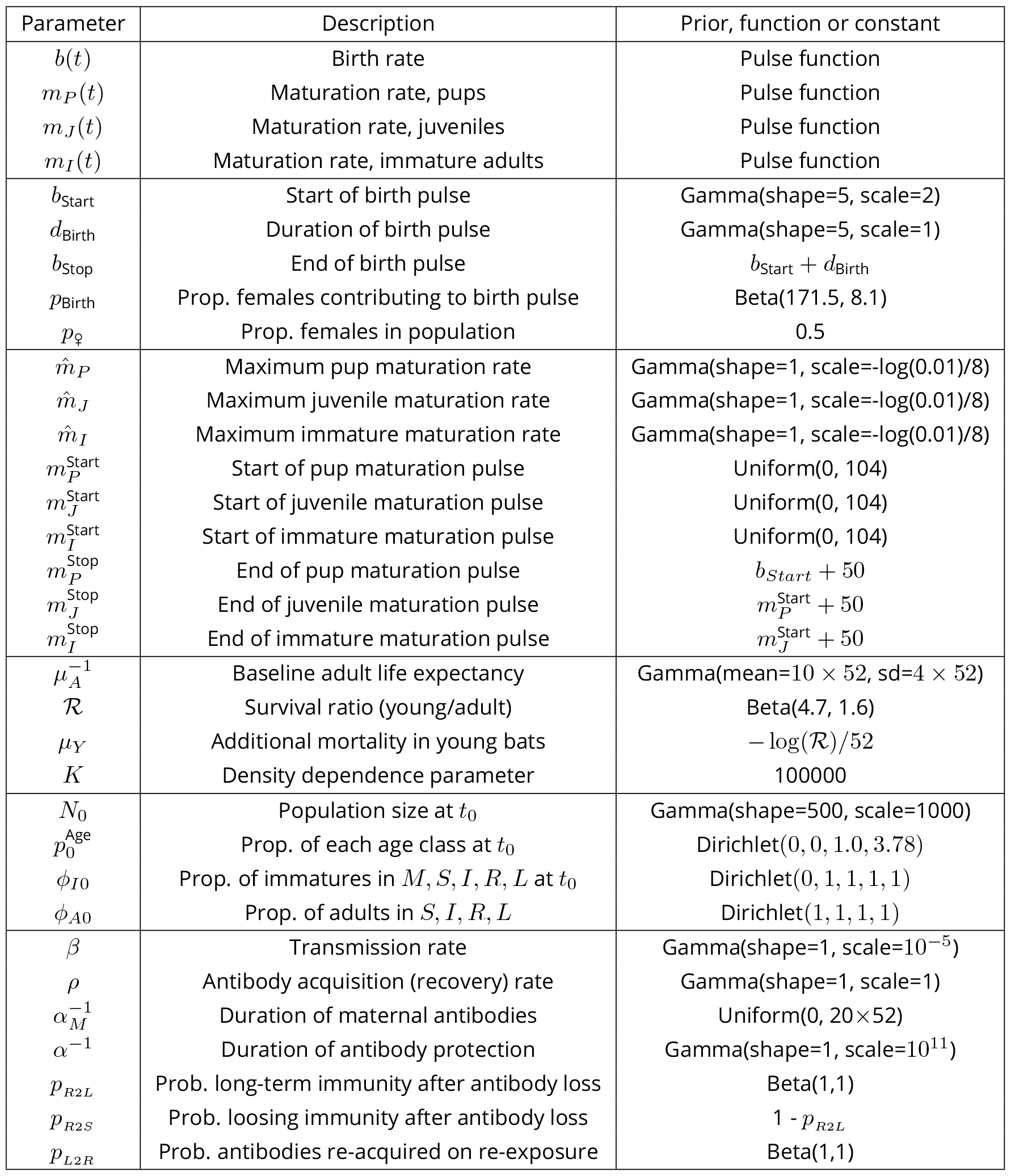
Parameters, priors and functions used for modelling the dynamics of Ebola virus circulation in the *Eidolon helvum* population of Yaounde, Cameroon (see Fig.1). These are presented in groups corresponding to: the four pulse functions; birth pulse parameters; maturation pulse parameters; mortality parameters; initial populations; and epidemiological parameters. The choice of priors is described in annex 2.

**Figure 1.**
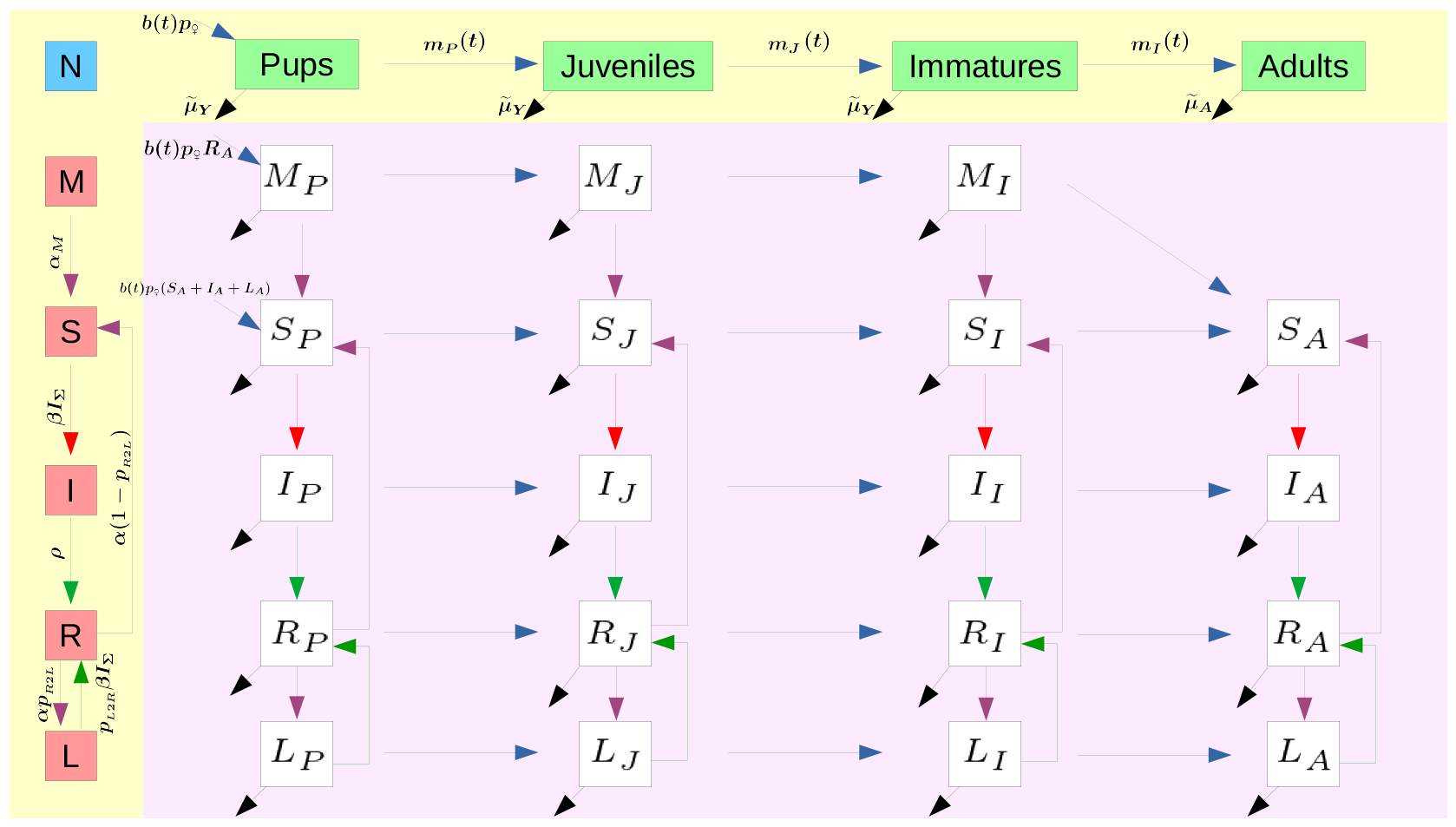
Schematic diagram of an age-structured MSIRL model used to analyse Ebola serology dynamics in *Eidolon helvum* from Yaounde, Cameroon. The total population *N* is divided into four age classes – pups (*P*), juveniles (*J*), immature adults (*I*) and adults (*A*) – and five epidemiological classes – maternal anti-bodies (*M*), susceptible (*S*), infected (*I*), recovered (*R*) and long-term immunity (*L*). Maturation through the age classes is controlled by a series of pulse functions (see annex 1), which lag behind a seasonal birth pulse. Density dependant mortality rates, 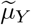 and 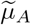, are specified for first-year and older individuals respectively. Anti-bodies are assumed detectable in individuals of the *M* and *R* compartments and undetectable for all other compartments. It is assumed that all pups from recovered mothers (*p*_♀_*R*_*A*_) start life with maternal antibodies (*M*_*P*_), and all other pups start life susceptible (*S*_*P*_).

**Figure 2.**
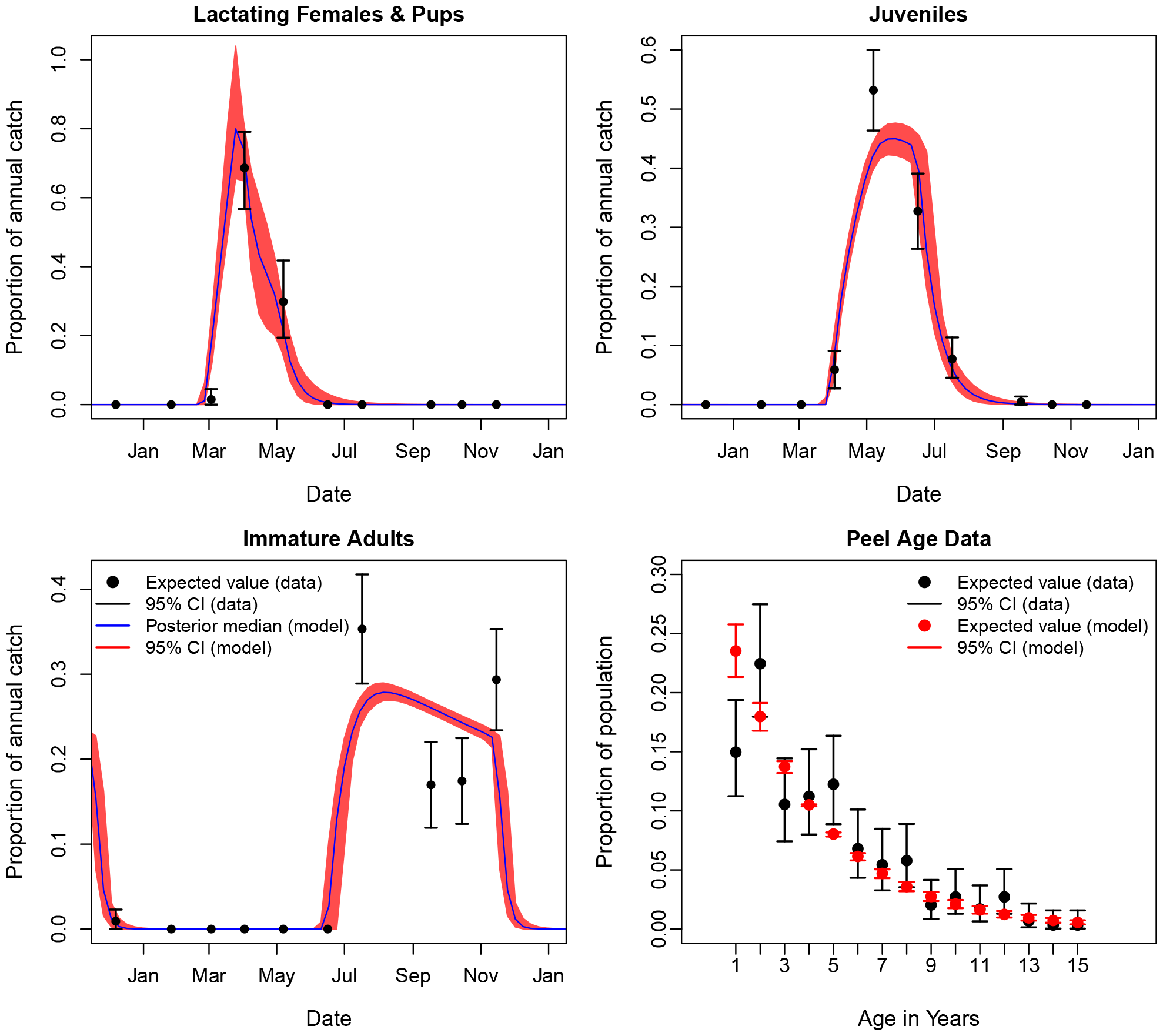
Comparison of model trajectories to age-structure data for *Eidolon helvum* in Yaounde, Cameroon (top row and bottom left), and of modelled adult survival to age estimates of adults based on tooth annuli data from Ghana (Peel, KS Baker, Hayman, Suu-Ire, et al., 2016).

The epidemiological model assumes that susceptible individuals become infected via a density dependent infection process with homogeneous mixing. Recovered individuals are assumed to transition to one of two classes – either they loose their immunity and return to being susceptible, or they enter a state of long-term immunity in which anti-bodies are not expressed unless they become re-exposed to the virus. Such a longterm immunity class has proved useful in modelling other bat-virus systems (Brook et al., 2019), and was included to avoid incompatibility between (i) rapid seroconversion among immature adults (and some juveniles), with seroprevalence reaching 60-80% in immatures, and (ii) a global seroprevalence of just 43% among all tested adults (figure 3). It was assumed that bats in the recovered and maternal antibody classes would express sufficient quantities of antibodies to test seropositive, whereas bats in all other epidemiological classes would test seronegative. We associated two parameters with the long-term immunity class: *p*_*R*2*L*_, the proportion of all individuals leaving the recovered class (i.e. loosing antibodies) that acquire long-term immunity, as opposed to loosing immunity and becoming susceptible again; and *p*_*L*2*R*_, the proportion of exposures to the virus that reinitialise anti-body production in bats with long-term immunity. Given a lack of evidence for vertical transmission for the related Marburg virus in Egyptian fruit bats (Towner et al., 2009), we assumed infectious females could only produce susceptible pups. We also assumed that the number of adult bats still expressing maternal antibodies was negligible, and thus omitted that category to reduce computation time.

**Figure 3.**
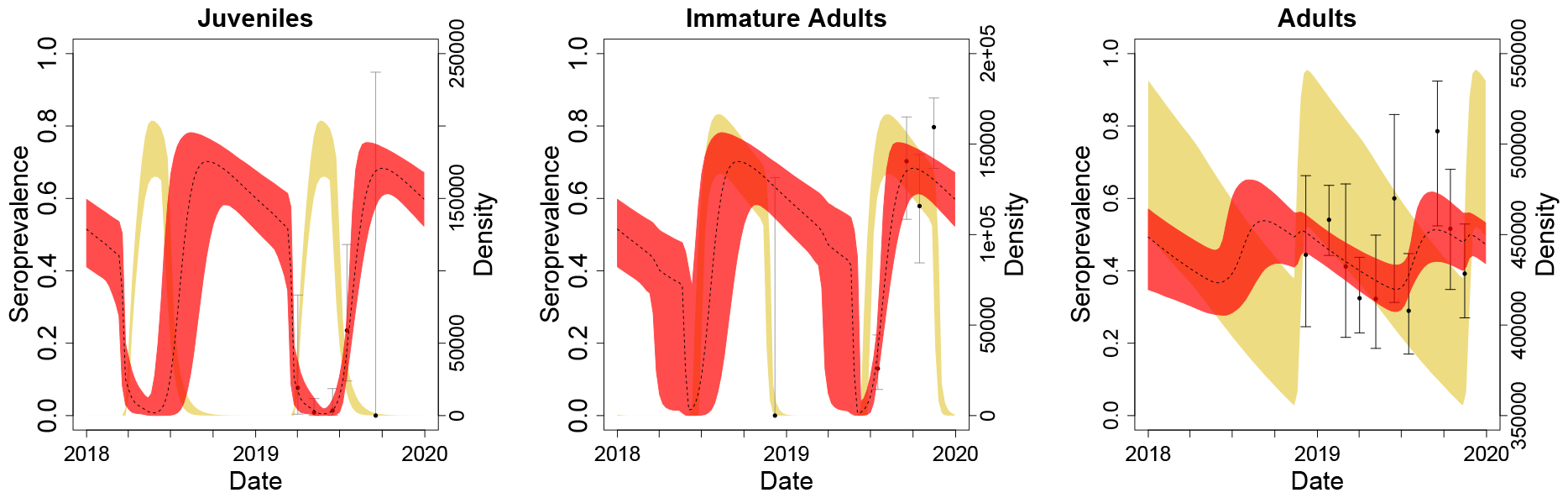
Seroprevalence data and estimates for 2018-2019. The mean and 95% credibility intervals for the seroprevalence data are shown as black dots and whiskers respectively. The median and 95% credibility intervals for the modelled seroprevalence are shown as dotted lines and red bands respectively. 95% credibility intervals for the density of individuals in each class are shown in beige.

Seasonal demographic dynamics were controlled via four pulse functions, which restrain when certain birth or maturation processes can or cannot occur. These functions are essentially smoothed (i.e. continuous) step functions that toggle whether or not a given step in the life cycle can be made at a given time. Each pulse function has three parameters: 1) the pulse start time; 2) the pulse end time; 3) and the rate at which individuals mature, or give birth, during the pulse. Nine of the twelve pulse function parameters were estimated as free parameters, whereas the three maturation pulses were constrained to end two weeks prior to the start date of the preceding pulse function of the following year (see table 2). This two week buffer ensured that individuals joining a given age class could not immediately mature to the following age class. A two week buffer size was chosen so that: 1) the buffer was large enough for overlap between the continuous pulse functions to be negligible; 2) each pulse function was wide enough so that only a negligible number of individuals remained in the age class when the maturation rate returned to zero. Further details of the pulse functions are provided in annex 1.

The transmission of Ebola virus within the *E. helvum* population of Yaounde, Cameroon, was modelled using the following system of ordinary differential equations (ODE):

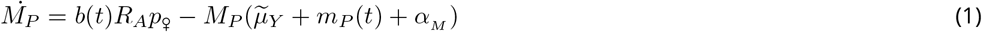

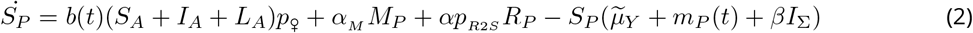

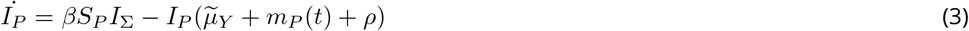

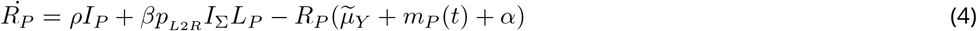

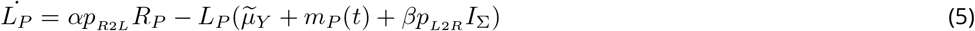

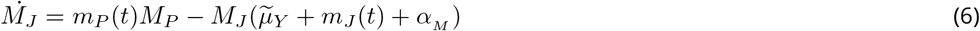

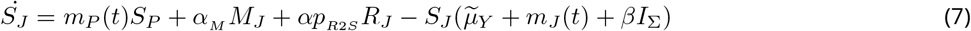

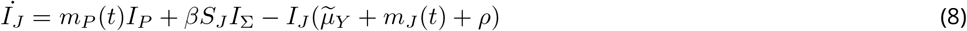

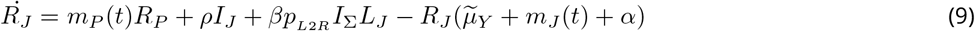

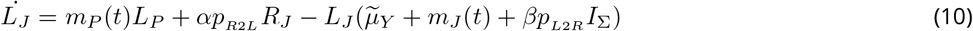

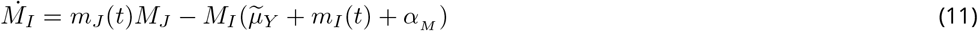

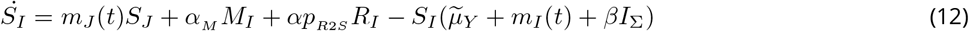

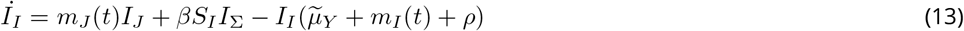

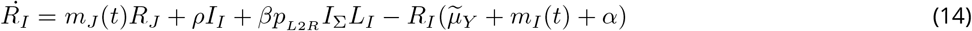

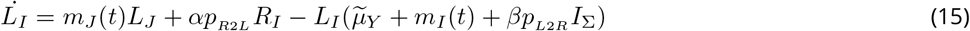

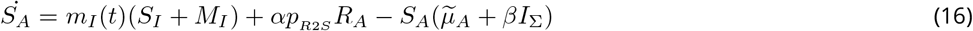

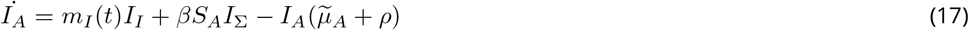

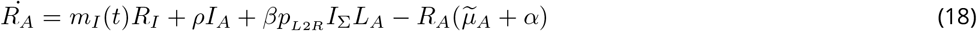

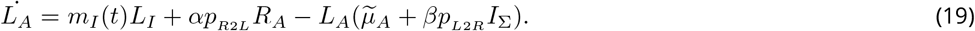

Table 2 provides a summary of model parameters and notation. Note, *I*_Σ_ = *I*_*P*_ + *I*_*J*_ + *I*_*I*_ + *I*_*A*_ is the total density of all infectious bats, and 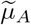 and 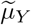 are density dependant mortality rates for adult and young bats respectively. Adult mortality was modelled as

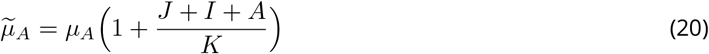

where *μ*_*A*_ is the mortality rate in the absence of competition, *K* is a density dependant parameter that contributes to determining the carrying capacity, and *J, I* and *A* provide the total population densities for juveniles, immatures and adults respectively. We assumed that, since pups and juveniles depend on their mothers, and that immature adults probably make mistakes that mature adults learn to avoid, then the mortality rates of non-adults should be equivalent to or higher than that of adults. Therefore, density dependant mortality among young bats was modelled as

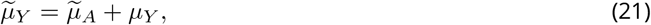

where *μ*_*Y*_ is the rate of additional mortality among young bats. Note, since *J, I* and *A* vary in time, so do 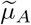 and 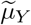. These density dependant mortality rates could therefore be represented using the notation 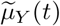 and 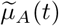, however, to simplify notation we adopt 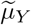 and 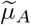 as shorthand alternative representations.

Let *𝒮*_*A*_(*t*) be a survival function that tracks how the survival probability of an adult bat decreases in continuous time. This survival function is described by the following differential equation

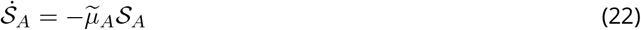

with the initial condition *𝒮*_*A*_(0) = 1. Let ***𝒮***_*A*_ and ***𝒮***_*Y*_ denote the annual survival probabilities for adult and young (<1 year) bats respectively ^1^. One way to obtain ***𝒮***_*A*_ would be to integrate equation 22 over a single 52 week year, i.e.

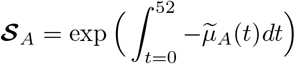

where 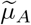(*t*) is the density dependant adult mortality (equation 20). Similar arguments for young bats give

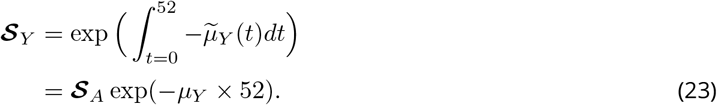

Thus, the additive nature of equation 21 permits us to parameterise *μ*_*Y*_ in terms of the ratio of the annual survival probabilities ***𝒮***_*Y*_ and ***𝒮***_*A*_. In other words

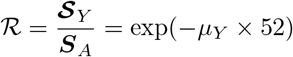

and

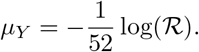

The advantage of this parameterization is that data were available for an informative prior on *ℛ* (see annex 2). In practice: integration of equation 22 was performed concurrently with the numerical integration of equations 1-19; ***𝒮***_*A*_ was obtained using equation 25; and ***𝒮***_*Y*_ was obtained using equation 23.

### Bayesian inference

A Bayesian approach was used to quantify uncertainty in model parameters, trajectories and derived metrics. Priors are detailed in table 2 and in annex 2. For each simulation of the ODE system performed during model fitting: state variables were initialized at the start of the year 2017; dynamics were simulated for three years; the ODE solver returned the state variables after each of 520 evenly spaced time steps per year; and the simulated trajectories were confronted with observed field data over the period December 2018 to November 2019. Starting the simulations in 2017 allowed a 23 month pre-data burn-in period in which the proportion of individuals in each category at time (*ϕ*_*It*_, *ϕ*_*At*_ and 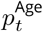) could converge from the wide range of possibilities permitted by the uninformative priors towards biologically plausible proportions driven by the model. The following subsections describe the various likelihood functions and penalties used for Bayesian inference, and outline how the model was used to address questions relating to (1) the mismatch between serology and PCR data, and (2) to the optimal timing of virology studies.

### Likelihood of age class data

The age distribution data (table 1) provides information as to when in the year we can expect to capture juveniles, immatures and lactating females – where the latter was used as a proxy for pups. It was suspected that between-class heterogeneity in capture rates could bias the absolute numbers of captures – therefore, the data were not used for calibrating between-class differences in density. Instead, we used this data to infer how the probability to capture a bat of a given class changes throughout the duration of the sampling period. Thus, for a given age class *j* ∈ {*P, J, I*}, the likelihood that the total number of captures were distributed across the various sampling dates as observed in the data was quantified assuming

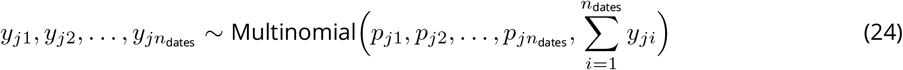

where *n*_dates_ is the number of observation dates, *y*_*ji*_ is the total number of bats of age class *j* captured at the *i*^th^ observation date and *p*_*ji*_ is the associated set of probabilities. The probabilities to sample a given pup (i.e. lactating female), immature or juvenile on the *i*^th^ sampling date were assumed to be proportional to the population density predicted by the system of ODEs at sampling time *t*_*i*_, thus,

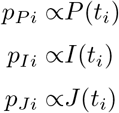

where 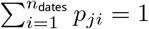 for any given age class *j*.

### Likelihood of tooth data

Tooth cementum annuli data (Peel, KS Baker, Hayman, Suu-Ire, et al., 2016) were used to inform estimates of adult mortality rates. Let *y*_*i*_ ∈ {1, 2, … } represent the age of bat *i* in years. A likelihood for a given bat’s age was obtained assuming

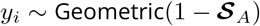

where ***𝒮***_*A*_ is the probability for an adult to survive one year. Following each simulation of three years, the annual adult survival probability was obtained as the ratio of the survival probabilities at the end of the final and penultimate years

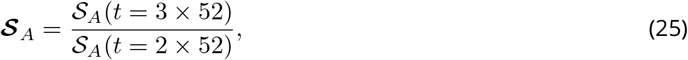

which is the conditional probability for an adult to survive the third year given that it survived the second year.

### Likelihood of serology data

The serology data (table 1) provides information about the number of seropositive individuals (*y*_*j*_(*t*)) of age class *j ∈ {J, I, A}* found in a sample of *n*_*j*_(*t*) individuals at time *t*. Thus, we assumed the following likelihood

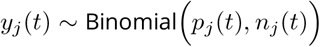

where

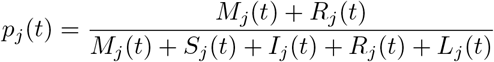

is the expected seroprevalence for age class *j* at time *t*.

### Penalties against demographic growth or decline

Due to an absence of longitudinal population census data, there were large uncertainties concerning the total population size at the beginning and end of the simulation period. We made the simplifying assumption that the *E. helvum* population was close to its carrying capacity and was approximately stable. Thus, we added penalty terms to the Bayesian model, to limit population growth or decline over the short simulation period and therefore constrain the potential distribution of starting population densities. These penalties were implemented as follows,

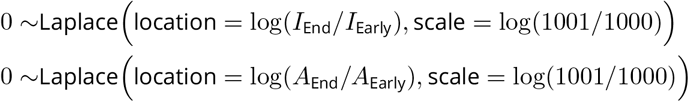

where *I*_Early_ and *I*_End_ are the total densities of immatures early on, and at the end of, a simulation, and *A*_Early_ and *A*_End_ are the total densities of adults early on, and at the end of, a simulation – where “Early” and “End” indicate the first model output for January following one year and three years of simulation respectively. The contribution to the total likelihood given by these penalties is greatest when there is zero population growth or decline over the last two years of the three year simulation period. The scale parameter controls the strength of the penalty. These penalties were only applied to the sizes of the adult and immature bat populations, because the other age classes were absent at the beginning of each year. Whilst it could arguably be reasonable to make the simplifying assumption that the total population size was roughly stable over the simulation period, a similar stability assumption for the epidemiological dynamics was considered to be too strong, since too little is known about the dynamics of Ebola in natural reservoirs – thus we did not use equivalent penalty terms to constrain the starting values of the various epidemiological compartments of the model.

### Markov chain Monte Carlo

Bayesian inference was based on Markov chain Monte Carlo sampling. An adaptive Metropolis Hastings block sampler was used to explore the posterior distribution of the model. Starting values for each parameter were based on the final values obtained from a previous short run of the algorithm. The sampler was run for 40 million iterations, with thinning set to 2000, and the first half of the samples were removed as a burn-in period. Thus, we obtained 10000 samples in total.

### Multi-annual cyclicity and skip years

An analysis of the long-term behaviour of the model was performed, with the aim of determining if seasonal patterns in prevalence were likely to be consistent (or not) from one year to the next. For each of the 10000 MCMC samples the ODE system was projected for 1100 years, with the first 1000 years removed as a burn-in period. The time vector sent to the ODE solver provided a temporal resolution of 10 steps per week. Each trajectory of infectious adults (*I*_*A*_) over the final 100 years was used to construct a recurrence plot (Marwan et al., 2007), using a threshold neighbourhood of 1 bat. In other words, each trajectory was used to construct a matrix with entries

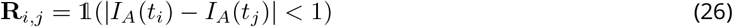

where 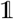 is the indicator function, *i* and *j* are indices for location along the time vector, and | *·* | represents the absolute value. Clearly, the main diagonal of any recurrence plot contains only ones (because *i* = *j* for each entry of the diagonal) and is uninteresting. However, any other diagonal containing only ones is interesting, because it informs about periodic (i.e. repeating) dynamics. Thus, we searched for the closest diagonal (to the principal diagonal) containing just ones, in order to identify *k*, the periodicity in years of any multi-annual pattern in *I*_*A*_. Thus 10000 values of *k* were tabulated in order to quantify uncertainty in the periodicity of the epidemiological dynamics. For this tabulation, we pooled all observations of 50 *< k <* 100 years, and 100 *< k*, to avoid potential false positives near the corners of the recurrence plots and to identify potentially chaotic trajectories.

For any simulation where we identified that *k >* 1 we searched for skip years, which we defined as any 52 week period within which the density of infectious adults (*I*_*A*_) consistently remains below one. Thus, when tabulating the various observed values of *k* we also tabulated the frequency of observing skip years as a function of *k*.

### Probability of not sampling an infectious bat

A key aim of this work was to quantify whether or not we should expect to see PCR positive bats in a typical sample given the fitted model of Ebola transmission in *E. helvum*. Let *𝒩*_*j*_(*t*) represent the sample size for bats of age class *j* obtained during a sampling campaign performed in week *t* – the probability to have zero infectious bats in this sample is:

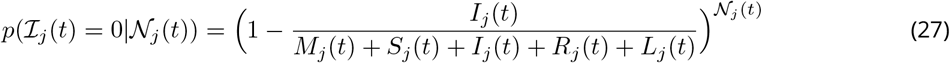

where *ℐ*_*j*_(*t*) is the number of infectious bats in the sample. We considered that 𝒩_*j*_(*t*) = 25 is a fairly typical scenario in a given sampling campaign, and thus plotted the evolution of *p*(ℐ_*j*_(*t*) = 0|𝒩_*j*_(*t*) = 25) in time for adult and immature bats, to provide an indication of when in the year would be an optimal time for sampling if viral extraction was the aim.

Similarly, we also calculated the probability of having not captured a single infectious bat given all the bats tested by PCR throughout the entire study,

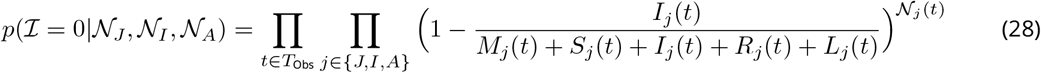

where *ℐ* is the total number of infectious bats sampled during the study, *𝒩*_*j*_ is the vector indicating how many bats of age class *j* were sampled in each sampling campaign, and *T*_Obs_ is the set of times for all of the observation campaigns.

### Implementation

All calculations were performed in R (R Core Team, 2022) version 4.2.1. Numerical integration of the ODE system (equations 1-19 and 22) was performed using the lsoda function in the deSolve package (Soetaert et al., 2010). Functions for the derivatives and Jacobian of the ODE system were coded in C. Bayesian inference was performed in `NIMBLE` (de Valpine, Paciorek, et al., 2022; de Valpine, Turek, et al., 2017), and the function nimbleRcall was used to call lsoda from inside `NIMBLE`. The package nimbleNoBounds (Pleydell, 2023) was used for improving the efficiency of adaptive Metropolis-Hastings sampling near the bounds of the parameter space. The R package CODA (Plummer et al., 2006) was used to perform convergence diagnostics on the MCMC output, and to provide the mean, median, 95% credibility interval and effective sample size (ESS) for each parameter. The effective sample size, which estimates the number of independent samples per parameter while accounting for auto-correlation, was calculated using the function `effectiveSize`. Whilst the system of ODEs was defined in continuous time, it is common for ODE solvers to discretize time – for each simulation lsoda was provided a time vector with intervals of 0.1 weeks to define when estimates for the state of the system were required. To economise on memory allocation we configured `NIMBLE` to store and use the state of the dynamic system at weekly time intervals.

## Results

### Inference from parameters

The posterior mean, median and 95% credibility intervals (shown in parentheses below) of each parameter, along with the annual survival and effective sample sizes (ESS) estimates obtained from 10000 MCMC samples, are presented in table 3. Twelve of the parameters were associated with ESS scores of 10000 or higher. The lowest ESS estimates were associated with the inverse of antibody loss rate (ESS(*α*^*−*1^) = 2289), the proportion of recovered individuals obtaining long-term immunity (ESS(*p*_R2L_) = 6024), and the proportion of adults with long-term immunity at the start of each simulation (ESS(*p*_0_(*L*|*Ad*)) = 7133).

**Table 3.** Summary of marginal posterior distributions for each parameter in the Ebola-*E*.*helvum* model. The mean, median and 95% credibility intervals for each parameter are presented, along with the effective sample size (ESS) estimated using the `effectiveSize` function of R package CODA.

| Parameter | Mean | 2.5% | Median | 97.5% | ESS |
| --- | --- | --- | --- | --- | --- |
| $b_{\text{Start}}$ | 8.04E+00 | 7.67E+00 | 8.04E+00 | 8.43E+00 | 10000 |
| $d_{\text{Birth}}$ | 9.12E+00 | 6.62E+00 | 9.31E+00 | 1.07E+01 | 7372 |
| $p_{\text{Birth}}$ | 9.54E-01 | 9.19E-01 | 9.56E-01 | 9.80E-01 | 10000 |
| $\hat{m}_P$ | 6.41E-01 | 3.31E-01 | 6.10E-01 | 1.11E+00 | 10000 |
| $\hat{m}_J$ | 4.58E-01 | 3.22E-01 | 4.37E-01 | 7.04E-01 | 10375 |
| $\hat{m}_I$ | 1.44E+00 | 7.71E-01 | 1.33E+00 | 2.75E+00 | 10000 |
| $m_P^{\text{Start}}$ | 1.25E+01 | 1.19E+01 | 1.25E+01 | 1.28E+01 | 10000 |
| $m_J^{\text{Start}}$ | 2.40E+01 | 2.30E+01 | 2.39E+01 | 2.55E+01 | 9381 |
| $m_I^{\text{Start}}$ | 4.58E+01 | 4.50E+01 | 4.57E+01 | 4.69E+01 | 11342 |
| $\mu_A^{-1}$ | 1.24E+03 | 1.09E+03 | 1.24E+03 | 1.41E+03 | 9532 |
| $\mathcal{R}$ | 5.12E-01 | 4.16E-01 | 5.10E-01 | 6.19E-01 | 9199 |
| $N_0$ | 4.92E+05 | 4.51E+05 | 4.92E+05 | 5.36E+05 | 9711 |
| $p_0(Im)$ | 2.09E-01 | 6.65E-03 | 1.67E-01 | 6.22E-01 | 10341 |
| $p_0(S\vert Im)$ | 2.43E-01 | 8.60E-03 | 1.99E-01 | 7.02E-01 | 10000 |
| $p_0(I\vert Im)$ | 2.60E-01 | 8.90E-03 | 2.17E-01 | 7.15E-01 | 9662 |
| $p_0(R\vert Im)$ | 2.45E-01 | 7.48E-03 | 2.00E-01 | 7.01E-01 | 10000 |
| $p_0(L\vert Im)$ | 2.52E-01 | 8.89E-03 | 2.07E-01 | 7.05E-01 | 10000 |
| $p_0(S\vert Ad)$ | 2.38E-01 | 6.09E-03 | 1.81E-01 | 7.18E-01 | 10000 |
| $p_0(I\vert Ad)$ | 2.76E-01 | 1.24E-02 | 2.41E-01 | 7.24E-01 | 9037 |
| $p_0(R\vert Ad)$ | 2.35E-01 | 7.17E-03 | 1.89E-01 | 6.91E-01 | 10000 |
| $p_0(L\vert Ad)$ | 2.51E-01 | 8.53E-03 | 2.07E-01 | 7.22E-01 | 7133 |
| $\beta$ | 5.57E-06 | 3.08E-06 | 4.96E-06 | 1.21E-05 | 7574 |
| $\rho$ | 6.77E-01 | 3.71E-01 | 6.06E-01 | 1.49E+00 | 7540 |
| $\alpha^{-1}$ | 7.51E+01 | 4.81E+01 | 6.96E+01 | 1.35E+02 | 2289 |
| $\alpha_M^{-1}$ | 1.11E+00 | 3.60E-01 | 1.04E+00 | 2.28E+00 | 9575 |
| $p_{R2L}$ | 6.33E-01 | 2.47E-01 | 6.54E-01 | 9.20E-01 | 6024 |
| $p_{L2R}$ | 1.70E-01 | 7.67E-03 | 1.47E-01 | 4.71E-01 | 9426 |
| $\mathcal{S}_Y$ | 3.91E-01 | 3.27E-01 | 3.90E-01 | 4.60E-01 | 9198 |
| $\mathcal{S}_A$ | 7.65E-01 | 7.42E-01 | 7.65E-01 | 7.87E-01 | 9474 |

The birth pulse is expected to start in the eighth week of the year and last nine (95% CI: 6.6 *−* 10.7) weeks. The three consequent maturation pulses are expected to start in weeks 12, 24 and 45 respectively. The ranges of the 95% credibility intervals for the four pulse function start times were (in chronological order) 0.76, 0.3, 1.5 and 1.9 weeks respectively. Annual survival probabilities were estimated as 39% (33% *−* 46%) and 76% (74% *−* 79%) in young and adult bats respectively. The estimated recovery rate, *ρ* = 0.67 (0.37 *−* 1.5), indicates that the expected duration of infections was 1.5 weeks (5 days – 19 days). Recovered bats are expected to produce antibodies for 75 (48 *−* 135) weeks, and maternal antibodies are expected to last 1.1 (0.36 *−* 2.3) weeks. The estimates of *p*_R2L_ indicate that roughly two thirds of recovered individuals pass to the long-term immunity class, although uncertainty was high (0.24 *−* 0.92). Only 17% of infectious attacks on individuals with long-term immunity re-initiate anti-body production, although uncertainty is large (0.7% *−* 47%).

A comparison of age-structure seasonality in the data and the model is presented in Fig. 2. The modelled trajectory of pup presence (red) closely follows the observed seasonal patterns in the number of lactating females (black). The model slightly underestimates the proportion of juveniles in May, but otherwise matches the juvenile data well - i.e. with overlap between the credibility intervals generated from the data and from the model. Seasonality in the presence/absence of immature adults is characterised well, although some considerable fluctuations in densities remain unexplained by the model. Similarly, there was considerable overlap between the credibility intervals calculated from the model and from the tooth-age data, albeit with some notable outliers among young adult bats.

### Seroprevalence dynamics

Comparisons of modelled and observed seroprevalence, in juvenile, immature adult and adult bats, are presented in Fig 3. The credibility intervals of observed and modelled seroprevalence overlap at all sampling dates. In juveniles and immature adults there is a large drop in seroprevalence when the maturation pulse functions permit the re-population of those age classes – by contrast, in adults there is a small increase in seroprevalence at the time when immatures start becoming adults. Seroprevalence increases in juveniles and immature adults during midsummer, with median seroprevalence rates being just 0.4% (0.05% - 1.8%) and 0.9% (0.007% - 3.8%) in weeks 22 and 24 (of 2019) respectively, and reaching 68% (60% - 74%) in week 40. Whilst this peak in seroprevalence is synchronised for the two classes, the density of juveniles is already reaching zero by that time, whereas the density of immature adults is reaching its maximum. A summertime upward trend is also observed in the seroprevalence of adults, with a median seroprevalence of 35% (29%-41%) in week 25 rising to 51% (43% - 62%) in week 38. The trajectories of both observed and modelled seroprevalence from early April to early May suggest that seroprevalence in juveniles drops considerably during this period – a continuation of the drop initiated one month earlier by the initiation of weaning in week 12.

### Period and predictability of long-term dynamics

The periods of multi-annual cyclicity in the dynamics of infectious adults (*I*_*A*_), identified using recurrence plots from 10000 simulations, are presented in table 4. Eighty nine percent of simulations resulted in dynamics with a period of one year – in these cases, the timing of the annual peak remained identical from one year to the next. Among the 11% of simulations which exhibiting more complex dynamics, 31% exhibited skip years. Nearly nine percent of simulations resulted in biennial (*k* = 2) cycles, 24% of which exhibited skip years. Sixty seven simulations resulted in four-year cycles, with 94% exhibiting skip years. Nineteen simulations exhibited *k >* 4 and *k <* 50. Seventy three simulations exhibited *k >* 50, with 48% exhibiting skip years. Examples of the types of trajectories possible under each value of *k* are presented in Fig. 4.

**Table 4.** Recurrence plot analysis results, providing the frequency distribution for various values of *k*, the period (in years) of dynamics in the density of infectious adults (*I*_*A*_), and the frequency of observing skip years in those patterns. Since recurrence plots were constructed from 100 year simulations, the maximum periodicity permitting at least one whole replication of a dynamic cycle was 50 years. Thus, we pool all simulations providing just partial evidence for periodicity in the 50-99 range. Similarly, we pool all simulations indicating *k ≥* 100, many of which are likely to have been chaotic. Four simulations resulted in extinction of the virus.

| Period $k$<br>(years) | Frequency | Frequency<br>with skips |
| --- | --- | --- |
| 1 | 8938 | 0 |
| 2 | 899 | 219 |
| 4 | 67 | 63 |
| 5 | 1 | 0 |
| 6 | 5 | 4 |
| 7 | 1 | 0 |
| 8 | 7 | 7 |
| 10 | 1 | 1 |
| 14 | 1 | 0 |
| 25 | 1 | 0 |
| 28 | 1 | 0 |
| 48 | 1 | 1 |
| 51-99 | 25 | 8 |
| $\geq 100$ | 48 | 27 |
| Extinct | 4 | N.A. |
| Total | 10000 | 330 |

**Figure 4.**
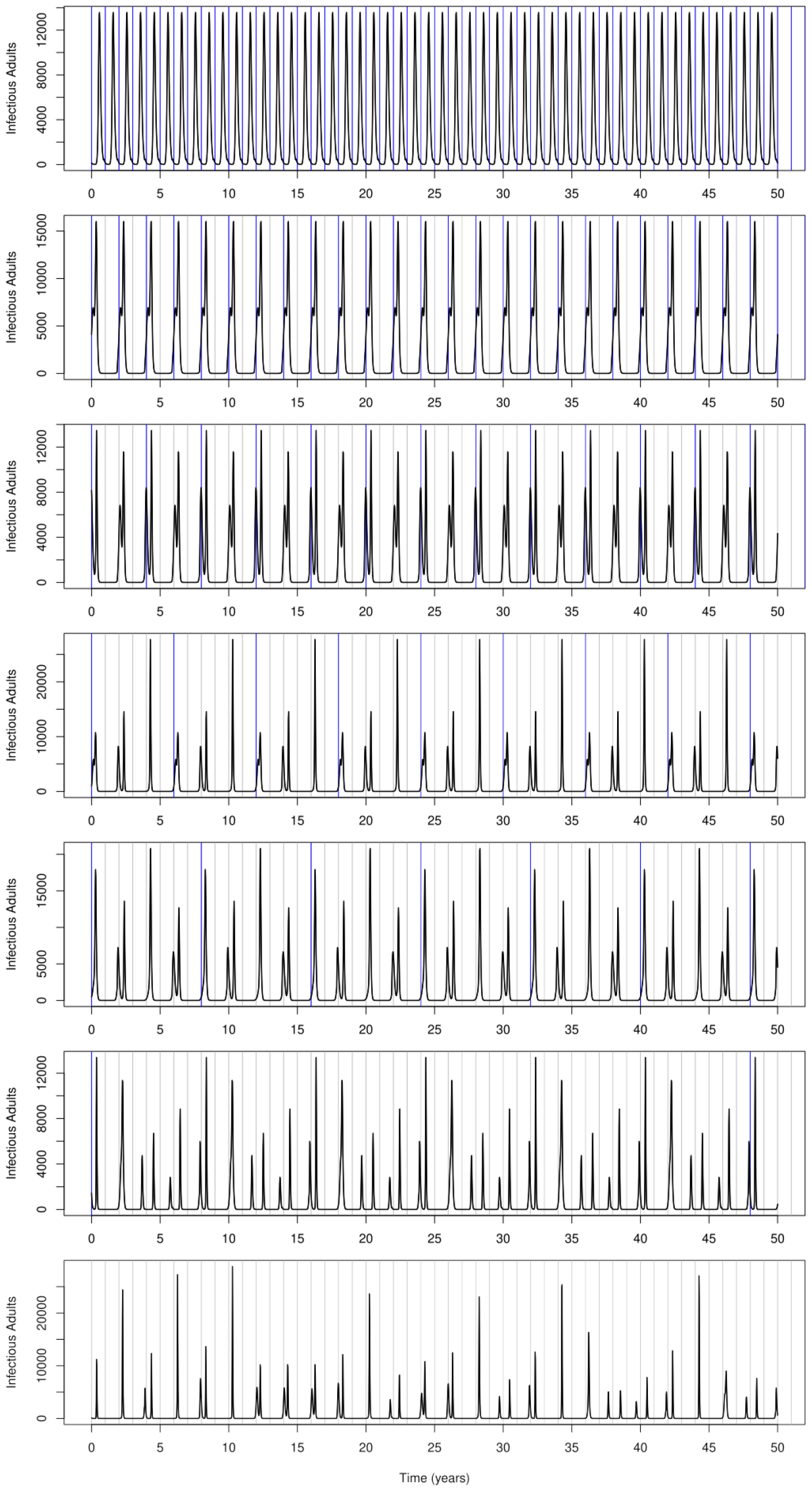
Example long-term trajectories of infectious adults under various values of period *k*. From top to bottom, *k* equals 1, 2, 4, 6, 8, 48 and *k >* 100 respectively. Vertical grey, and blue, lines depict the start of each year, and the period *k*, respectively. Skip years (any 52 week period without an outbreak) are evident in several examples. A higher (than one bat) threshold in the recurrence plot definition (equation 26) could clearly result in *k* = 8 and not *k* = 48 in the sixth example. The dynamics in the final example appear to be chaotic.

The timing of the annual peak in infectious adults, and the relation ship between that timing and the size of the peak, is presented in Fig. 5. An annual peak in the density of infectious adults is most likely in weeks 30 - 31 (p=0.63), in weeks 17 - 27 (p=0.05), or in weeks 48 - 52 (p=0.045). Weeks 21, 25 and 27 are associated with the greatest expected outbreak sizes (24610, 24411 and 24528 infectious adults respectively), despite bi-modality in outbreak size during that period of the year. The expected outbreak size in weeks 30 and 50 are 8128 and 9618 respectively.

**Figure 5.**
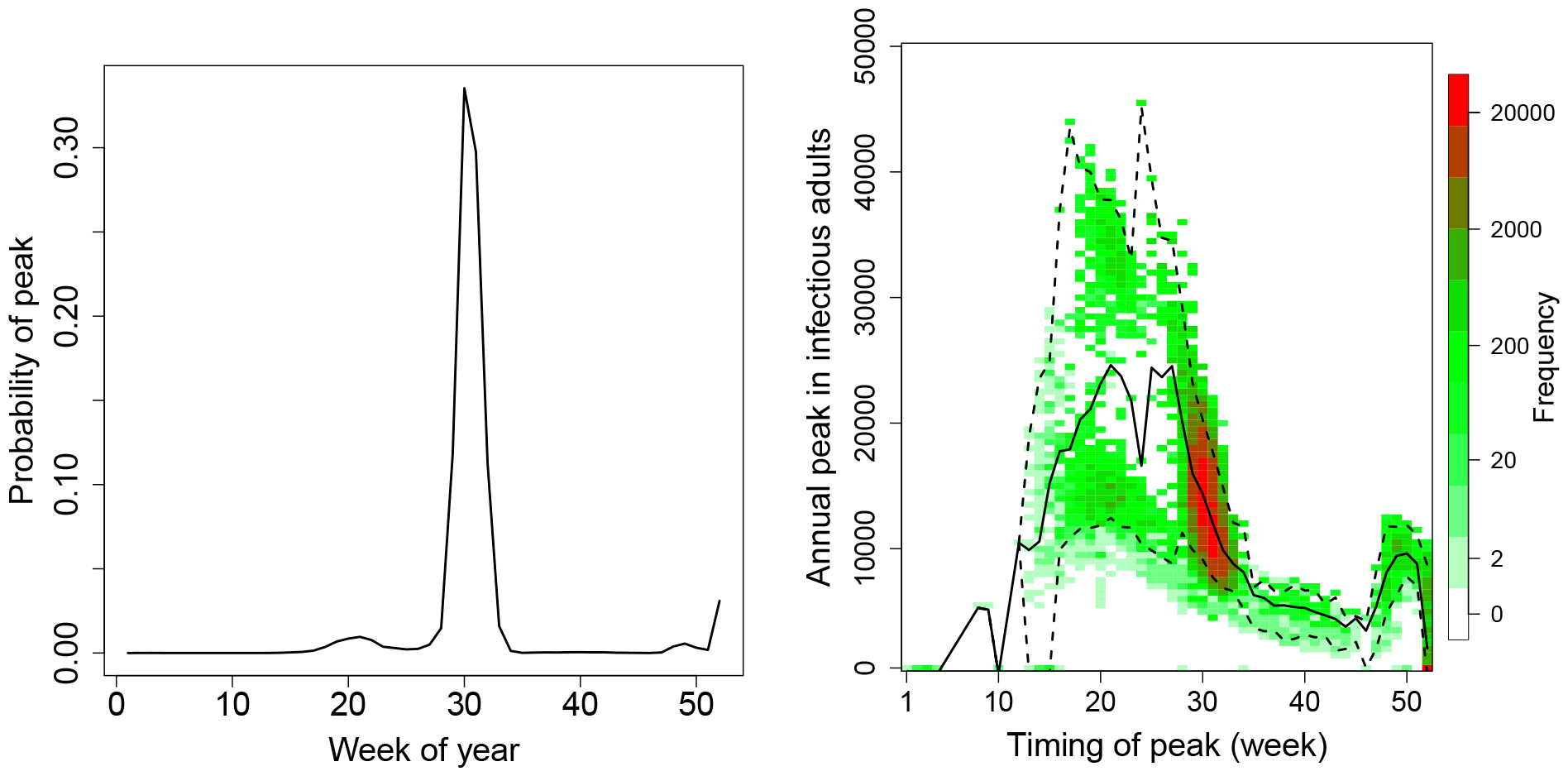
Seasonal trend in the probability that the annual peak in infectious adults falls within a given week (left), and the relationship between the timing and size of outbreak peaks (right). These results are based on the final 100 years of 10000 simulations, each 1100 years in length. The probability of the peak given the week of the year was derived as the expected value derived from these one million years of simulated output (black line, left plot). The frequency distribution for the timing and size of peak *I*_*A*_ density for each of these one million years of simulated output is represented via a white-green-red colour scale (right plot), with the weekly mean and 95% credibility intervals represented as solid and dashed lines respectively.

### Probability of not sampling an infectious bat

Seasonality across 2019 in the probabilities to not have an infectious individual in a sample of 25 adults and 25 young bats are represented graphically in figure 6. The expected values of these probabilities were minimised in week 31 in both young and adult bats, and were 0.02 (95% CI: 0.0039 – 0.069) and 0.48 (0.29 – 0.70) respectively.

**Figure 6.**
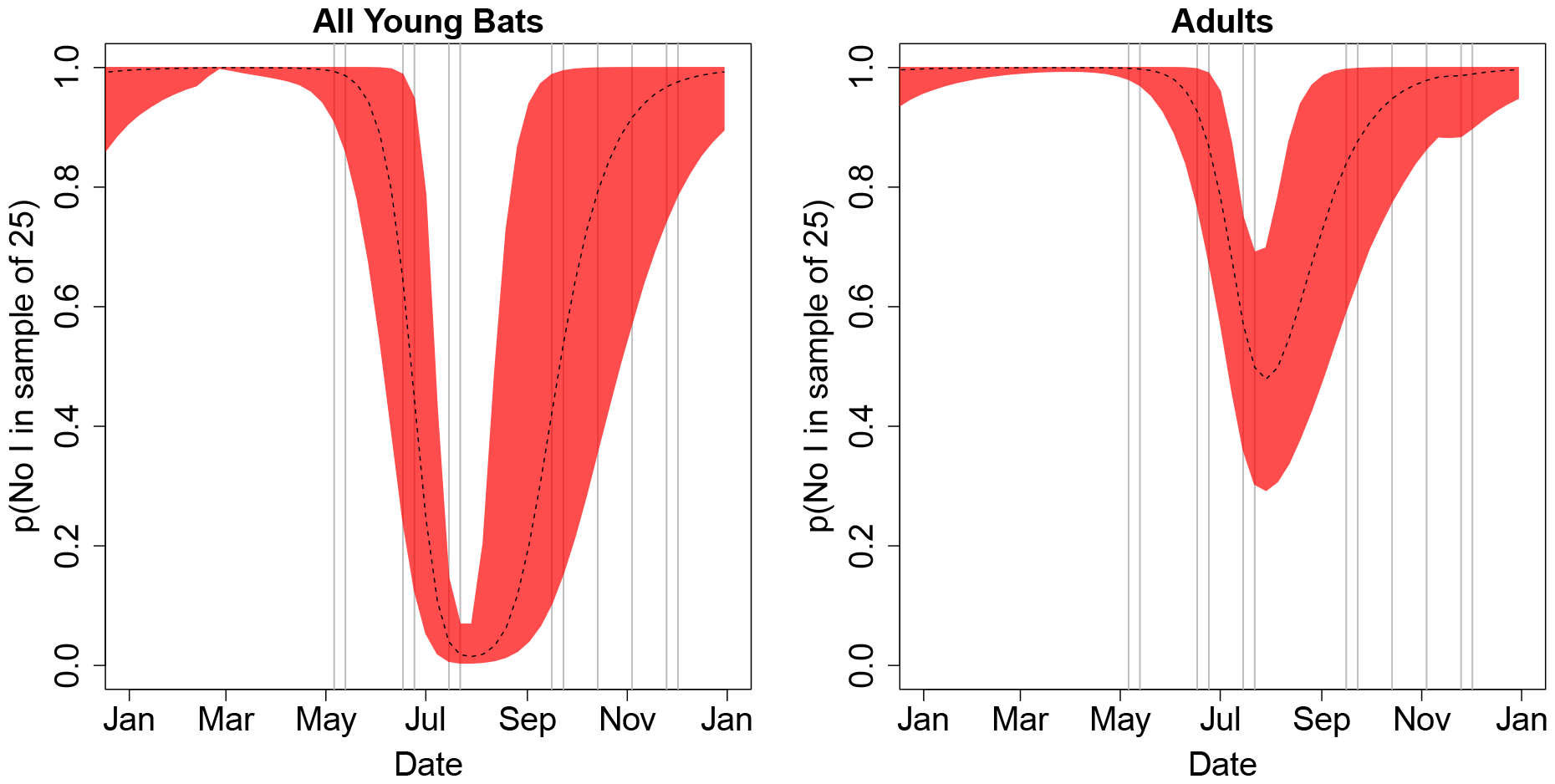
Probability of not capturing an infectious bat in a sample of 25 pre-adult (left) or adult (right) bats, across 2019. Grey vertical lines indicate weeks at which bats were captured for PCR analysis, with a mean sample size of 25.

The probability to not have an infectious bat in the samples tested by PCR in the (Djomsi et al., 2022) study was 0.00052 (6.6 *×* 10^*−*9^ – 4.2 *×* 10^*−*3^). These probabilities are greatest during the first four to five months of the year. Uncertainty in these probabilities is greatest in late summer and early autumn.

## Discussion

### Model overview and fit

The current work presents a Bayesian analysis of an age-structured epidemiological model of Ebolavirus transmission in *Eidolon helvum*. The model simulates both demographic and epidemiological dynamics, and was calibrated to ecological and serological data collected previously in Cameroon (Djomsi et al., 2022) and age structure data from Ghana (Peel, KS Baker, Hayman, Suu-Ire, et al., 2016). A key component of the model is a series of four seasonally dependant pulse functions, which control when females can produce pups, and when maturation between successive age classes can occur. Uncertainty in the estimated starting times of those pulse functions was low, with the 95% credibility interval being less than two weeks wide in all four cases. Some outliers in the age-structure data were observed (Fig. 2) and are likely linked to neglected ecological mechanisms, such as heterogeneity in dispersion patterns, food availability and survival. Nevertheless, the model trajectories provide a succinct summary of trends observed in both the age-structure and serology data – the most notable trend being the sharp increase in seroprevalence in late summer (Fig. 3).

### Inference from parameter estimates

Our analyses indicate that, on average, 76% of adults and 39% of young bats survive each year. Infections are expected to last one and a half weeks. Maternal antibodies are expected to provide protection for just 1.1 weeks on average, thus the annual birthing pulse leads rapidly to growth in the pool of susceptible individuals, which in turn typically leads to increased transmission and seasonal outbreaks. Somewhat similar patterns of maternal antibody loss, followed by acquisition as young adults, and with seroprevalence in adults stabilising at roughly 60%, have also been reported for Lagos bat virus and henipavirus in *E. helvum* (Peel, KS Baker, Hayman, Broder, et al., 2018) – however, the mean duration of protection from maternal antibodies in that study was estimated to be half a year. Following experimental infections with canine distemper virus in adult female *Pteropus hypomelanus* and natural infections of Hendra virus in adult female *Pteropus alecto* serological tests could still detect maternal antibodies in pups of up to 7.5 and 8.5 months of age (Epstein, ML Baker, et al., 2013). The duration of protection from maternal antibodies estimated in the current study does appear to be low compared to estimates from other studies for other viruses, however, uncertainty was low (despite a very uninformative prior) which suggests that the result was really driven by the observed data. Further studies could be useful to verify why maternal antibodies play an apparently less important role here, compared to other host-virus systems.

Here, the expected duration of antibodies in recovered bats was estimated to be 75 weeks, or possibly as much as 135 weeks – although again, this duration of projection from antibodies is shorter than the four years and twelve years estimated for henipavirus and Lagos bat virus respectively in Peel, KS Baker, Hayman, Broder, et al. (2018). Given the shorter half-life of detectable antibodies in the adult bats of the current study, it is perhaps less surprising that maternal immunity appears to be shorter here than in other studies.

The posterior distribution of *p*_R2L_ indicated that the majority of individuals loosing antibodies are expected to enter a form of long-term immunity – and 97.5% of samples indicated *p*_R2L_ *>* 0.24, thus some form of long-term immunity appears to be likely. This result replicates modelling results of Brook et al. (2019), who described a similar phenomenon in henipavirus transmission in *Eidolon dupreanum, Pteropus rufus* and *Rousettus madagascariensis* fruit bats in Madagascar. Moreover, experimental infections suggest that Egyptian Fruit Bats (*Rousettus aegyptiacus*) continue to exhibit long-term protection to Marburg virus 17-24 months after an original infection despite waning expression of virus-specific IgG antibodies (Schuh, Amman, TK Sealy, et al., 2017).

### Probability of serology-PCR mismatch

A key aim of the current work was to explore an apparent mismatch between seroprevalence data – which suggest Ebola-related virus circulation in juvenile and sexually immature bats – and the results of PCR tests – which have failed to detect a positive sample among the 456 oral and rectal swabs tested from 366 bats (152 juveniles and 214 immature adults)(Djomsi et al., 2022). Here the probability to not have an infectious bat among all the samples tested by PCR was estimated to be 0.00052 (95% CI: 6.6 *×* 10^*−*9^ - 4.2 *×* 10^*−*3^), which confirms a paradoxical mismatch between the results of the serology and PCR tests.

The circulation pattern observed in the serology data, and replicated in our model, is apparently driven by seasonal pulses of young susceptible bats entering the population, fueling an annual resurgence of viral circulation, and playing a key role in viral persistence. That birthing patterns play an important role for contributing to the timing of outbreaks has been reported for various other host-pathogen systems (Cappelle, Furey, et al., 2021; Jolles et al., 2021; Mariën et al., 2020; Peel, Pulliam, et al., 2014), which supports the argument that the seasonal patterns observed in the serology data really are linked to viral circulation. However, in the absence of confirmed positive control samples for ebolaviruses in bats the calibration of a serological test is challenging, therefore there is a risk that a low cut-off value could have inflated the frequency of false positive results. Indeed, Djomsi et al. (2022) tried several methods to identify a cut-off value – however, even the most stringent of those cut-offs suggested the presence of bats that were seropositive to ebolaviruses and seasonality in transmission. Cross-reactivity between different ebolaviruses has been documented in humans (Diallo et al., 2021) and in experimentally infected *Rousettus aegyptiacus*, where limited cross reactivity with other filoviruses was also documented (Schuh, Amman, TS Sealy, et al., 2019). Such results suggest that the serological signal observed in that study did come from the circulation of Ebola-related viruses and not other filoviruses. Nevertheless, false positive reactivity with other pathogens cannot be excluded for the serological assay used in our study, which may explain why all PCR tests remained negative – i.e. the viruses actually circulating and causing positive serology in *E. helvum* might not be in the detection range of the panfilovirus PCR of Djomsi et al. (2022). However, other factors could also explain the lack of positive PCR test results, even if Ebola-related viruses actually are circulating within the bat population.

One alternative possibility is that low sensitivity of the PCR assay may have lead to many false negative test results and may therefore explain the mismatch between the serological and PCR data. PCR assays designed to detect viral families may have lower sensitivity than PCR targeting specific viruses. For example a Bomabalivirus-specific real-time PCR assay detected an addiational postivie sample than the filovirus ‘family level’ cPCR assay used by Goldstein et al. (2018).

Furthermore, samples taken from infectious sylvatic bats are likely to have very low viral loads compared to experimentally infected bats or sick naturally infected humans for whom the PCR assays have been designed. If PCR sensitivity is an issue, then developing a more sensitive PCR should help, so long as it is not associated with a decrease in specificity. Indeed, if unknown Ebola-related viruses are actually circulating in the population, designing a specific PCR assay would prove challenging. Moreover, future studies that succeeded to identify or isolate those viruses would greatly clarify the epidemiological picture.

Finally, another potential explanation for the negative PCR results, despite the apparent circulation of Ebola-related viruses, may be the absence of viral excretion in the rectal and oral swab samples collected. During an experimental inoculation of *Rousettus aegyptiacus* with Ebola virus, none of 36 swab samples taken 3-10 days post infection tested positive by PCR, although Ebola RNA was detected in the blood of one bat and the lungs and liver of another (Paweska et al., 2016). Transmission routes other than the fecal-oral or oral-oral routes may be involved in the transmission of Ebola-related viruses in *E. helvum*. In rare cases Ebola virus has been detected in various samples from humans, and a sexual route of transmission has been demonstrated (Christie et al., 2015; Mate et al., 2015; Thorson et al., 2016). The large majority of samples taken from bats so far have been oral and rectal swabs. Taking multiple samples from bats, including organs may help to clarify this point. Ethical questions would arise from such a protocol involving bat euthanasia, and the balance between improving our understanding of the ecology of ebolaviruses and animal well-being should be discussed by ethics experts.

### Complex dynamics and optimal timing for sampling

A key aim of the current study was to predict the optimal timing for identifying or isolating the virus(es) responsible for the sero-conversions observed in *E. helvum*. Clearly, when planning field sampling schemes, it can be highly beneficial to have as complete an understanding as possible concerning the complexity of viral dynamics in a sylvatic host population. Some bat-borne zoonotic viruses are known to exhibit complex multi-year inter-epizootic periods, which have been attributed to interactions between population density changes, waning immunity, and viral recrudescence (Cappelle, Hoem, et al., 2020; Epstein, Anthony, et al., 2020). Results from our long-term simulations indicate a degree of uncertainty regarding whether or not complex multi-annual dynamics in the number of infectious bats are to be expected. Ten percent of our simulations suggest that the period of cyclicity could be greater or equal to two years, and 31% of that subset of simulations suggest that there may be periods of twelve months or more where prevalence rates remain close to zero. Such “skip years” are a well known phenomenon in mathematical epidemiology (Stone et al., 2007; Subramanian et al., 2020; Zhao et al., 2018) and arise when the size of the susceptible population remains below a threshold required for an outbreak for prolonged periods of time. Clearly, whether or not skip years occur is an important question for field-virologists interested in sampling sylvatic hosts for virus isolation. Here the probability that the system exhibits skip years was estimated as 0.033, which is low but not completely negligible either.

Actually, almost 90% of our long-term simulations suggested that the dynamics of ebolavirus in *E. helvum* in Cameroon may be relatively simple. The most likely scenario appears to be: one outbreak occurs per year; the size of those outbreaks is somewhat consistent; and the peak of each outbreak likely occurs during weeks 30 and 31 of the year (p=0.63). Thus, a sampling campaign centered at these dates would most likely be optimal. However, our uncertainty analysis does not eliminate the possibility of more complex patterns where the peak in the number of infectious bats could occur at any time after the first three months of the year, and where the size and timing of outbreaks are related. Given this uncertainty in the timing and size of outbreaks, it could also be worth sampling in weeks 17-27, because although the probability to have an outbreak in this period is lower, the size of outbreaks predicted in this period can be greater. Any outbreaks occurring after week 35 would only generate low prevalence rates, thus it could be challenging to isolate the virus during this period. These results can be used to target periods when ebolavirus circulation can be expected to be greatest, and to help optimise the sample sizes required to have a high probability of sampling at least one infectious bat – which can help limit the number of bats euthanized for the purpose of viral isolation.

### Limitations and future research

Various limitations should be kept in mind when interpreting the results presented in the current work. For example, our modelling neglects: stochasticity in population dynamics, transmission and recrudescence (Muñoz et al., 2022; Peel, Pulliam, et al., 2014); spatial dynamics and migration (Richter and Cumming, 2006); between-year variation in the timing and success of birth pulses (Adole et al., 2016); potential long-term carriers (Forrester, 2018); temporal changes in environmental stress that may affect susceptibility (Lafferty and Holt, 2003); and age-dependant heterogeneity in contact rates (Rohani et al., 2010). Future modelling studies should consider using sensitivity analysis to assess whether or not neglecting such mechanisms can have important consequences on the long-term trajectories of disease transmission and on the optimal timing of sampling. Moreover, the current work has focused on one host, one serological test and is based on just over one year of field data. We cannot eliminate the possibility that multiple Ebola-related viruses contributed to the observed trends in serology, because of a lack of specificity of the serological tests. The limitations of this study highlight the importance of conducting long-term field monitoring, for the calibration of models, assessing their predictions and for fully elucidating the complex dynamics of Ebola-related viruses in sylvatic host communities.

## Conclusions

The current paper presents modelling work that addresses a paradoxical observation in straw coloured fruit bats, where young bats exhibit rapid seroconversion for ebolavirus antibodies whilst confirmation by PCR remains elusive. The probability of this contradictory observation is estimated to be one in two thousand. The potential causes of this mismatch have been discussed and remain the focus of future research. This work provides novel insights in to the nature of the seasonality of ebolavirus transmission in fruit bats and provides predictions which can help with the design of future field programs for isolating circulating Ebola viruses.

## Acknowledgements

We thank the staff of CREMER (Joseph Moudindo, Aime Mebanga, Thomas Atemkeng, Eitel Mpoudi Ngole) for logistical support in the field in Cameroon.

## Data, script and code availability

Scripts and data used in the current analysis are available online at https://doi.org/10.5281/zenodo.8276172.

## Supplementary information

Details regarding the pulse functions used to control seasonality are provided in annex 1. Details regarding the parameterisation of priors for three parameters are provided in annex 2.

## Annex 1 Pulse functions

Seasonal flow of individual bats through the four-class life cycle model was controlled via a series of four pulse functions. The scaled product of two logistic curves was used to define a single pulse, and modulo arithmetic was used so that this pulse could be applied to an unlimited number of years. Thus, the rate of a given life-cycle process (i.e. birth, or maturation) was modelled as a function of time *t* as follows:

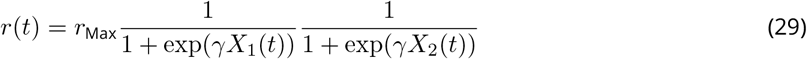

With

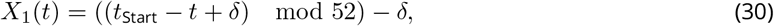

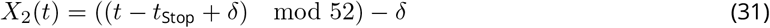

and

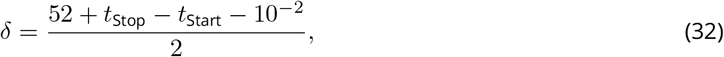

where *t*_Start_ gives the start of the pulse (i.e. the centrality parameter for the first logistic curve), *t*_Stop_ gives the end of the pulse (i.e. the centrality parameter for the second logistic curve), modulo arithmetic permits the recycling of the pulse function over multiple years, *δ* provides a shift that eliminates artefacts arising from edge effects under most biologically reasonable combinations of parameters, and *γ* is a shape parameter controlling how rapidly the rate *r*(*t*) passes from zero to *r*_Max_, and back again. In practice we fix *γ* at ten. Note, *t*_Stop_ *> t*_Start_.

For the birth pulse function, *r*_Max_ represents the within-season birth rate, which we note as *b*_Max_, which we define as

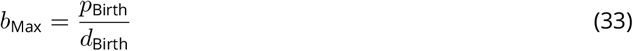

where *d*_Birth_ is the duration of the birth pulse, and *p*_Birth_ is the proportion of females expected to give birth (to a single pup) during the birth pulse.

## Annex 2: Parameterisation of priors

Prior distributions for all fitted parameters are summarised in table 2 of the main text. In the following subsections we outline our choices for how the prior distributions for each of the estimated model parameters were specified. Typically our choices for prior distributions were classic. For example: the beta distribution was used to model scalar proportions; Dirichlet distributions were used for vectors of proportions that sum to one; and gamma distributions were used as priors for positive scalars such as rates or sejourn times. Recall, an exponential distribution is a gamma distribution with a shape parameter of one.

### Prior for *b*_Start_ and *d*_Birth_

For the start and duration of the birth pulse (*b*_Start_ and *d*_Birth_ respectively) semi-informative priors were chosen to represent the knowledge and uncertainties of ecologists familiar with the *E*.*helvum* population of Yaounde.

For the start of the birth pulse, *b*_Start_, a gamma distribution was chosen with an expected value of 10 and a standard deviation of approximately 4.5. This provides a distribution with approximately 90% of its mass distributed between the 4th and 18th week of the year.

For the duration of the birth pulse, *d*_Birth_, a gamma distribution was chosen with an expected value of 5 and a standard deviation of approximately 2.2, providing a distribution with approximately 90% of its mass distributed between 2 and 9 weeks.

### Prior for *p*_Birth_

The proportion of females giving birth each year, *p*_Birth_, was modelled using data from Hayman, McCrea, et al. (2012) and a beta-binomial model, which is a natural choice for modelling proportions with binary data. According to that paper, the expected value and 95% confidence interval of *p*_Birth_ are 0.96 and (0.92,0.98) respectively. We sought to identify the parameters of a beta distribution that would minimise the *L*^2^ norm of errors between fitted values and these three data points. Using the optim function in R, we identified that

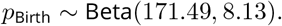

For further details, see the script hayman.R.

### Prior for 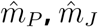 and 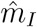

The maximum maturation rates of pups, juveniles and immatures (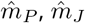 and 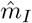) were given exponential prior distributions, with the scale parameter set so that an expected 99% of individuals completed the given stage of the life cycle within eight weeks. The expected time for 99% of individuals to complete the life stage becomes 76 weeks (or 3.5 weeks) if the maturation rate was set to the 10^*th*^ (or 90^*th*^) percentile of its prior distribution (respectively) – suggesting that this the prior is only weakly informative within a biologically plausible range of values.

### Prior for 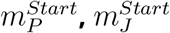 and 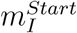

The start of the pulse functions controlling maturation (to subsequent life stages) of pups, juveniles and immatures were given uniform priors. The bounds on those uniform distributions were set to 0 and 104 weeks, making them uninformative over the expected development time of *E. helvum*.

### Prior for 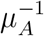

The baseline mortality rate for adults, *μ*_*A*_, is the mortality rate that is expected in the absence of competition with other bats. Thus, it is the expected mortality rate when bat densities are close to zero. We set a mildly informative prior on its inverse, the baseline life expectancy. For that, we used a gamma distribution with an expected value of ten years and a standard deviation of four years. For this distribution 99% of the mass corresponds to the expected life expectancy being less than 21.6 years.

### Prior for *ℛ*

According to Hayman, McCrea, et al. (2012) the expected annual survival probability and 95% confidence intervals is 0.63 and (0.27, 0.88) for adult bats. Using arguments similar to the previous section on *p*_Birth_, we used optim to minimise the *L*^2^ norm of the errors between these three data points and fitted values, giving the following model of adult survival

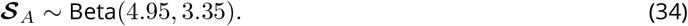

Similarly, Hayman, McCrea, et al. (2012) reported the expected annual survival probability and 95% confidence intervals for young bats are 0.43 and (0.16, 0.77) respectively. To ensure that ***𝒮***_*Y*_ *<* ***𝒮***_*A*_, we assumed ***𝒮***_*Y*_ = *R****𝒮***_*A*_ and that *R* could be modelled using a beta distribution. Thus, we sought to identify parameters for *R* that could minimise an *L*^2^ norm between the three data points and their equivalent “fitted values”. We used Monte Carlo approximation to obtain these “fitted values” as follows.

Assume the following model for *ℛ*,

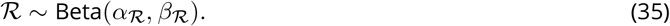

For a given set of parameters (*α*_*ℛ*_, *β*_*ℛ*_), we simulated 10001 values from equations 34 and 35. Those vectors were multiplied to obtain 10001 samples of ***𝒮***_*Y*_, and kernel density estimation was applied to these samples to obtain an empirical distribution for ***𝒮***_*Y*_. This empirical distribution was used to identify the fitted expected value and 95% credibility interval, which were then used to calculate the *L*^2^ norm. Minimising the *L*^2^ norm resulted in obtaining the following prior

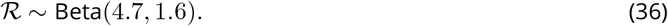

For further details, see our script hayman.R.

### Prior for *N*_0_

The total population size at the start of each simulation (*N*_0_) is a parameter that cannot be known with precision, given the lack of census data or capture-mark-recapture studies. However, experience in the field indicates that Yaounde’s *E. helvum* population is extremely large, and probably consists of several hundreds of thousands of individuals. We adopted a prior that was informative about the order of magnitude of the population - representing uncertainty in the total population size via a gamma distribution, with an expected value of 5 *×* 10^5^, a standard deviation of 2.2 *×* 10^4^ and 2.5^*th*^ and 97.5^*th*^ percentiles of 4.6 *×* 10^5^ and 5.4 *×* 10^5^ respectively. The purpose of this prior was to constrain *N*_0_ within a likely order of magnitude, in order to facilitate the estimation of the other parameters.

### Prior for 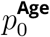

Before simulating dynamics with an epidemiological model, it is necessary to set the initial conditions of the system, i.e. the state of each compartment at time zero. For an age structured model, this includes setting the initial population sizes for each age class. We do that by parameterizing in terms of the total population size at time zero, *N*_0_, and the proportion of that population associated with each age class, 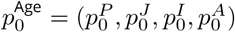. Since pups and juveniles are absent at the start of the year we set their initial proportions to zero. The prior on *N*_0_ was set to approximate the unknown population size in Yaounde. Thus, we simply needed to set a prior for the proportion of immatures, 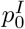, and it’s compliment, 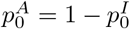. We outline how we did that here.

Since 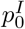 is a probability, it was natural to assign a beta distribution and to seek data on which to base the hyper-parameters. Assuming that the annual adult survival is constant with age, the population age structure follows a geometric series, and the proportion of individuals of a given age relative to all individuals of the same age or greater is constant. Thus, we used tooth cementum annuli data (Peel, KS Baker, Hayman, Suu-Ire, et al., 2016) to estimate that proportion, and thereby obtain an estimate for the proportion of immature adults in the population at the start of the year, just prior to the spring birth pulse. For each age *t*, in years, we modelled the proportion of bats of age *t* among all bats of age *t* or more as a beta-binomial model with uniform prior, i.e.

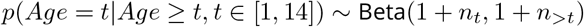

where *n*_*t*_ is the number of sampled bats of age *t, n*_*>t*_ is the number of sampled bats older than *t* and *t* is any integer in the interval [1, 14] – since the oldest bat in the data set was 15 years old. A weighted average of these 14 priors was calculated to obtain a general prior

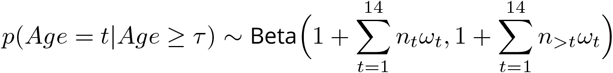

where the weights 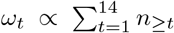 account for the diminishing sample size as bats die each year. Since *p*(*Age* = *t*|*Age ≥ τ*) is constant under the constant mortality assumption, it might therefore be reasonable to use it as a prior for 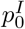. This procedure resulted in the prior

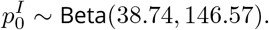

However, in practice, this prior lead to mismatches with data that suggested that very few bats were still being classified as immature at the start of the year (fig. 2). Thus, we maintained the expected value of this prior, but relaxed the variance so to not exclude zero as the proportion of immatures at the start of January. This relaxation resulted in the following prior

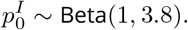

The density functions of these priors, and the 14 distributions used to build them, are shown graphically in figure 7.

**Figure 7.**
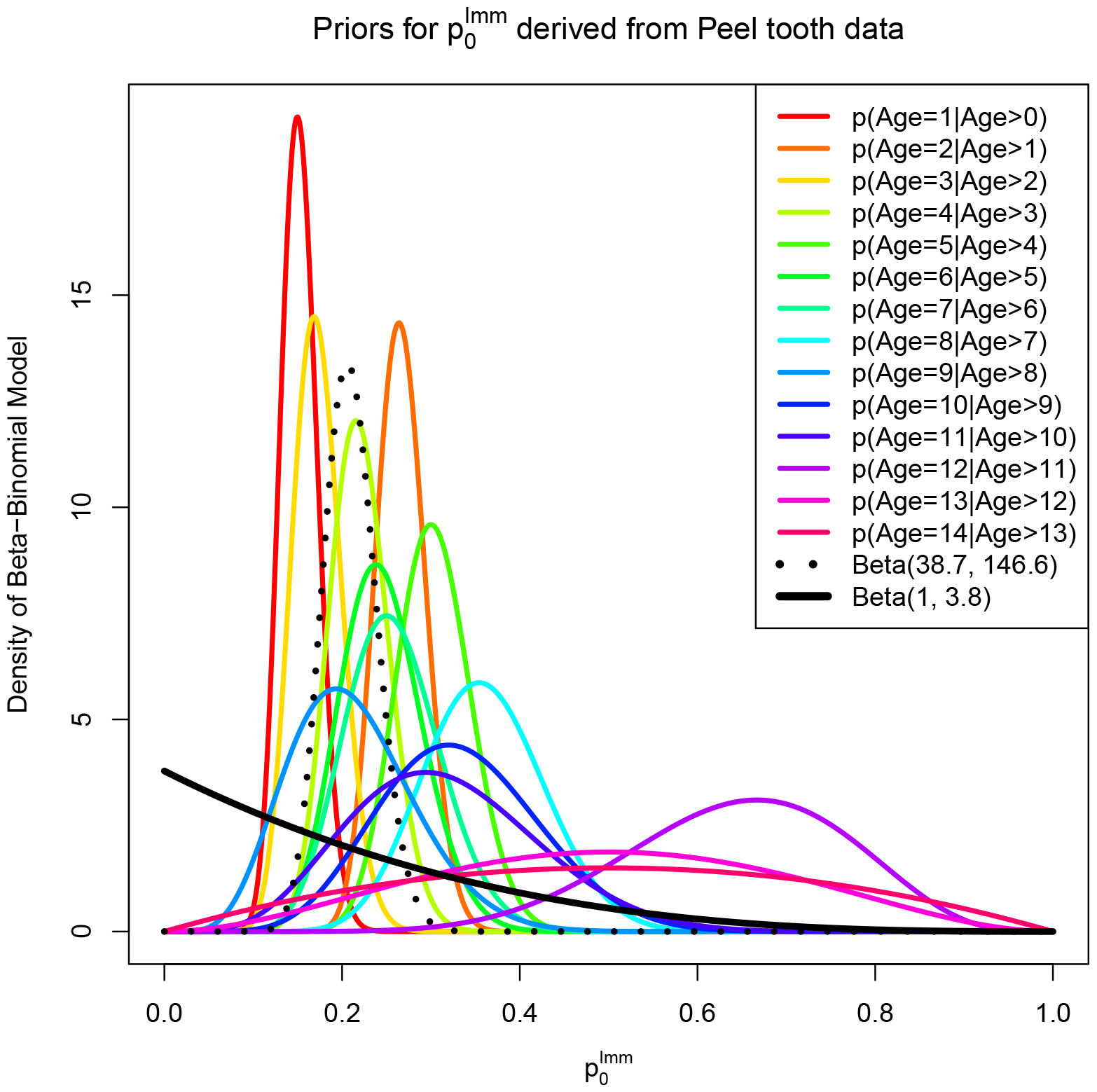
Posterior distributions for beta-binomial models of the proportion of a given age of adult in years (*t*) in the sub-population of bats the same age or more. Tooth cementum annuli data (Peel, KS Baker, Hayman, Suu-Ire, et al., 2016) were used to calculate these distributions, fixing *t* at integer values in the interval [1, 14]. The weighted average of those 14 distributions (dotted line) proved to be overly restrictive as a prior. So the variance of the prior was relaxed to not exclude zero (black line), providing a prior for the proportion of immature adults in the *E*.*helvum* population at the start of the year.

### Prior for *ϕ*_*I*0_ and *ϕ*_*A*0_

The proportions of immatures and adults within each of the epidemiological classes at the start (*t* = 0) of each simulation (*ϕ*_*I*0_ and *ϕ*_*A*0_ respectively) were given Dirichlet priors. These priors were parameterised to be uninformative, with the exception that we assumed no bats in either of these age classes will carry maternal antibodies at the start of the year.

### Prior for *β*

Experience with our model indicated that uninformative priors for the transmission rate *β* do not work well. Thus, it was important to restrain *β* from being so large that the posterior distributions became biologically implausible. To do this, we assumed an exponential (or equivalently, a gamma distribution with shape equal to one) prior, to penalise against very large values of *β*. To obtain a reasonable expected value for this prior we asked roughly how many new infections might a single infectious individual generate in one week when introduced into a completely susceptible population (and neglecting all other transitions). In other words, we asked what might be a roughly reasonable value for the product *βSI*, where *I* = 1 and *S* = *E*[*N*_0_]. Since *E*[*N*_0_] = 5 *×* 10^5^ we opted for *E*[*β*] = 10^*−*5^ so that *a priori* the expected number of secondary cases in one week is five. Thus we used the following prior

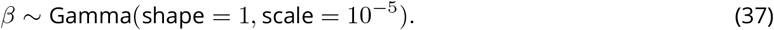

Setting *β* to the 1st or 99th percentile of this prior leads to the product *βSI* being 0.05 or 23 respectively.

### Prior for 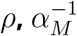 and *α*^*−*1^

For the recover rate, we set an exponential prior with an expected value of one week – a value consistent with many virus infections in humans. In other words we assumed that

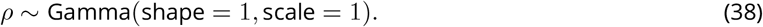

For the expected duration of maternal antibodies (inverse of antibody loss rate) we set an uninformative uniform prior over the range of zero to twenty years

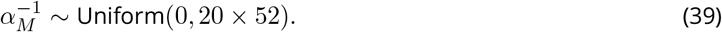

For the expected duration of maintaining antibodies following infection, we specified the following non-informative exponential prior

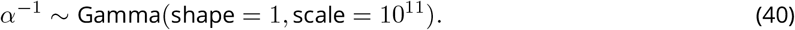

### Prior for *p*_*R*2*L*_ and *p*_*L*2*R*_

The probabilities of developing long-term immunity following the loss of antibodies, *p*_*R*2*L*_, and of re-acquiring antibodies when exposed to the virus whilst in a state of long term immunity, *p*_*L*2*R*_, were assigned uniform uninformative priors. In other words,

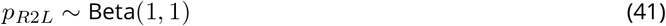

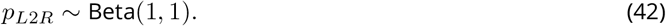

## Footnotes

1 Note, different fonts are used for susceptible adults (*S*_*A*_), adult survival (*S*_*A*_) and the annual adult survival probability (***S***_*A*_).

## References

Adole T, J Dash, and PM Atkinson (2016). A systematic review of vegetation phenology in Africa. Ecological Informatics 34, 117–128.

Brook CE, HC Ranaivoson, CC Broder, AA Cunningham, J. Héraud, AJ Peel, L Gibson, JL Wood, CJ Metcalf, and AP Dobson (2019). Disentangling serology to elucidate henipa-and filovirus transmission in Madagascar fruit bats. Journal of Animal Ecology 88, 1001–1016.

Cappelle J, N Furey, T Hoem, TP Ou, T Lim, V Hul, O Heng, V Chevalier, P Dussart, and V Duong (2021). Longitudinal monitoring in Cambodia suggests higher circulation of alpha and betacoronaviruses in juvenile and immature bats of three species. Scientific Reports 11, 24145.

Cappelle J, T Hoem, V Hul, N Furey, K Nguon, S Prigent, L Dupon, S Ken, C Neung, V Hok, et al. (2020). Nipah virus circulation at human–bat interfaces, Cambodia. Bulletin of the World Health Organization 98, 539.

Caron A, M Bourgarel, J Cappelle, F Liégeois, HM De Nys, and F Roger (2018). Ebola virus maintenance: if not (only) bats, what else? Viruses 10, 549.

Champagne M, J Cappelle, A Caron, T Pouliquen, A Samoura, MI Doumbouya, G Thaurignac, AK Ayouba--Ahidjoans--Keita, M Peeters, M Bourgarel, and H De--Nys (Submitted). Effects of biological and environmental factors on filovirus serology in bats in Guinea.

Christie A, GJ Davies-Wayne, T Cordier-Lasalle, DJ Blackley, AS Laney, D. Williams, SA Shinde, M Badio, T Lo, SE Mate, et al. (2015). Possible sexual transmission of Ebola virus—Liberia, 2015. Morbidity and Mortality Weekly Report 64, 479.

De Nys HM, PM Kingebeni, AK Keita, C Butel, G Thaurignac, CJ Villabona-Arenas, T Lemarcis, M Geraerts, N Vidal, A Esteban, et al. (2018). Survey of Ebola viruses in frugivorous and insectivorous bats in Guinea, Cameroon, and the Democratic Republic of the Congo, 2015–2017. Emerging infectious diseases 24, 2228.

de Valpine P, C Paciorek, D Turek, N Michaud, C Anderson-Bergman, F Obermeyer, C Wehrhahn--Cortes, A Ro-dríguez, D Temple--Lang, S Paganin, and J Hug (2022). NIMBLE: MCMC, Particle Filtering, and Programmable Hierarchical Modeling. Version 0.13.1. 10.5281/zenodo.1211190.

de Valpine P, D Turek, C Paciorek, C Anderson-Bergman, D Temple Lang, and R Bodik (2017). Programming with models: writing statistical algorithms for general model structures with NIMBLE. Journal of Computational and Graphical Statistics 26, 403–417. 10.1080/10618600.2016.1172487.

Diallo MSK, A Ayouba, G Thaurignac, MS Sow, C Kpamou, TA Barry, P Msellati, JF Etard, M Peeters, R Ecochard, et al. (2021). Temporal evolution of the humoral antibody response after Ebola virus disease in Guinea: a 60-month observational prospective cohort study. The Lancet Microbe 2, e676–e684.

Djomsi DM, FA Mba Djonzo, I Ndong Bass, M Champagne, A Lacroix, G Thaurignac, A Esteban, HM De Nys, M Bourgarel, JF Akoachere, et al. (2022). Dynamics of antibodies to Ebolaviruses in an Eidolon helvum bat colony in Cameroon. Viruses 14, 560.

Epstein JH, SJ Anthony, A Islam, AM Kilpatrick, S Ali Khan, MD Balkey, N Ross, I Smith, C Zambrana-Torrelio, Y Tao, et al. (2020). Nipah virus dynamics in bats and implications for spillover to humans. Proceedings of the National Academy of Sciences 117, 29190–29201.

Epstein JH, ML Baker, C Zambrana-Torrelio, D Middleton, JA Barr, E DuBovi, V Boyd, B Pope, S Todd, G Crameri, et al. (2013). Duration of maternal antibodies against canine distemper virus and Hendra virus in pteropid bats. PLoS one 8, e67584.

Feldmann H, A Sprecher, and TW Geisbert (2020). Ebola. New England Journal of Medicine 382, 1832–1842.

Forrester JV (2018). Ebola virus and persistent chronic infection: when does replication cease? Annals of Trans-lational Medicine 6.

George DB, CT Webb, ML Farnsworth, T. O’Shea, RA Bowen, DL Smith, TR Stanley, L. Ellison, and CE Rupprecht (2011). Host and viral ecology determine bat rabies seasonality and maintenance. Proceedings of the National Academy of Sciences 108, 10208–10213.

Glennon EE, DJ Becker, AJ Peel, R Garnier, RD Suu-Ire, L Gibson, DT Hayman, JL Wood, AA Cunningham, RK Plowright, et al. (2019). What is stirring in the reservoir? Modelling mechanisms of henipavirus circulation in fruit bat hosts. Philosophical Transactions of the Royal Society B 374, 20190021.

Goldstein T, SJ Anthony, A Gbakima, BH Bird, J Bangura, A Tremeau-Bravard, MN Belaganahalli, HL Wells, JK Dhanota, E Liang, et al. (2018). The discovery of Bombali virus adds further support for bats as hosts of ebolaviruses. Nature microbiology 3, 1084–1089.

Hayman DT (2015). Biannual birth pulses allow filoviruses to persist in bat populations. Proceedings of the Royal Society B: Biological Sciences 282, 20142591.

Hayman DT, AD Luis, O Restif, KS Baker, AR Fooks, C Leach, DL Horton, R Suu-Ire, AA Cunningham, JL Wood, et al. (2018). Maternal antibody and the maintenance of a lyssavirus in populations of seasonally breeding African bats. PloS one 13, e0198563.

Hayman DT, R McCrea, O Restif, R Suu-Ire, AR Fooks, JL Wood, AA Cunningham, and JM Rowcliffe (2012). Demography of straw-colored fruit bats in Ghana. Journal of mammalogy 93, 1393–1404.

Hayman DT, M Yu, G Crameri, LF Wang, R Suu-Ire, JL Wood, and AA Cunningham (2012). Ebola virus antibodies in fruit bats, Ghana, West Africa. Emerging infectious diseases 18, 1207.

Hranac CR, JC Marshall, A Monadjem, and DT Hayman (2019). Predicting Ebola virus disease risk and the role of African bat birthing. Epidemics 29, 100366.

Jacob ST, I Crozier, WA Fischer, A Hewlett, CS Kraft, MAdL Vega, MJ Soka, V Wahl, A Griffiths, L Bollinger, et al. (2020). Ebola virus disease. Nature reviews Disease primers 6, 1–31.

Jolles A, E Gorsich, S Gubbins, B Beechler, P Buss, N Juleff, LM de Klerk-Lorist, F Maree, E Perez-Martin, OL Van Schalkwyk, et al. (2021). Endemic persistence of a highly contagious pathogen: Foot-and-mouth disease in its wildlife host. Science 374, 104–109.

Lafferty KD and RD Holt (2003). How should environmental stress affect the population dynamics of disease? Ecology Letters 6, 654–664.

Leroy EM, B Kumulungui, X Pourrut, P Rouquet, A Hassanin, P Yaba, A Délicat, JT Paweska, JP Gonzalez, and R Swanepoel (2005). Fruit bats as reservoirs of Ebola virus. Nature 438, 575–576.

Letko M, SN Seifert, KJ Olival, RK Plowright, and VJ Munster (2020). Bat-borne virus diversity, spillover and emergence. Nature Reviews Microbiology 18, 461–471.

Mariën J, B Borremans, C Verhaeren, L Kirkpatrick, S Gryseels, J Goüy--de--Bellocq, S Günther, CA Sabuni, AW Mas-sawe, J Reijniers, et al. (2020). Density dependence and persistence of Morogoro arenavirus transmission in a fluctuating population of its reservoir host. Journal of Animal Ecology 89, 506–518.

Marwan N, MC Romano, M Thiel, and J Kurths (2007). Recurrence plots for the analysis of complex systems. Physics reports 438, 237–329.

Mate SE, JR Kugelman, TG Nyenswah, JT Ladner, MR Wiley, T Cordier-Lassalle, A Christie, GP Schroth, SM Gross, GJ Davies-Wayne, et al. (2015). Molecular evidence of sexual transmission of Ebola virus. New England Journal of Medicine 373, 2448–2454.

Muñoz F, DRJ Pleydell, and F Jori (2022). A combination of probabilistic and mechanistic approaches for predicting the spread of African swine fever on Merry Island. Epidemics 40, 100596.

Munster VJ, DG Bausch, E De--Wit, R Fischer, G Kobinger, C Muñoz-Fontela, SH Olson, SN Seifert, A Sprecher, F Ntoumi, et al. (2018). Outbreaks in a rapidly changing Central Africa—lessons from Ebola. New England Journal of Medicine 379, 1198–1201.

Olival KJ and DT Hayman (2014). Filoviruses in bats: current knowledge and future directions. Viruses 6, 1759– 1788.

Paweska JT, N Storm, AA Grobbelaar, W Markotter, A Kemp, and P Jansen--van--Vuren (2016). Experimental inoculation of Egyptian fruit bats (Rousettus aegyptiacus) with Ebola virus. Viruses 8, 29.

Peel AJ, KS Baker, DT Hayman, CC Broder, AA Cunningham, AR Fooks, R Garnier, JL Wood, and O Restif (2018). Support for viral persistence in bats from age-specific serology and models of maternal immunity. Scientific reports 8, 1–11.

Peel AJ, KS Baker, DT Hayman, R Suu-Ire, AC Breed, GC Gembu, T Lembo, A Fernández-Loras, DR Sargan, AR Fooks, et al. (2016). Bat trait, genetic and pathogen data from large-scale investigations of African fruit bats, Eidolon helvum. Scientific data 3, 1–11.

Peel AJ, J Pulliam, A Luis, R Plowright, T O’Shea, D Hayman, J Wood, C Webb, and O Restif (2014). The effect of seasonal birth pulses on pathogen persistence in wild mammal populations. Proceedings of the Royal Society B: Biological Sciences 281, 20132962.

Pleydell DRJ (2023). nimbleNoBounds: Transformed Distributions for Improved MCMC Efficiency. Version 1.0.2. 10.5281/zenodo.6399163.

Plummer M, N Best, K Cowles, and K Vines (2006). CODA: Convergence Diagnosis and Output Analysis for MCMC. R News 6, 7–11.

R Core Team (2022). R: A Language and Environment for Statistical Computing. R Foundation for Statistical Computing. Vienna, Austria.

Richter H and G Cumming (2006). Food availability and annual migration of the straw-colored fruit bat (Eidolon helvum). Journal of Zoology 268, 35–44.

Rohani P, X Zhong, and AA King (2010). Contact network structure explains the changing epidemiology of pertussis. Science 330, 982–985.

Schuh AJ, BR Amman, TK Sealy, JR Spengler, S. Nichol, and JS Towner (2017). Egyptian rousette bats maintain long-term protective immunity against Marburg virus infection despite diminished antibody levels. Scientific reports 7, 8763.

Schuh AJ, BR Amman, TS Sealy, TD Flietstra, JC Guito, S. Nichol, and JS Towner (2019). Comparative analysis of serologic cross-reactivity using convalescent sera from filovirus-experimentally infected fruit bats. Scientific Reports 9, 1–12.

Soetaert K, T Petzoldt, and RW Setzer (2010). Solving Differential Equations in R: Package deSolve. Journal of Statistical Software 33, 1–25. 10.18637/jss.v033.i09.

Stone L, R Olinky, and A Huppert (2007). Seasonal dynamics of recurrent epidemics. Nature 446, 533–536.

Subramanian R, V Romeo-Aznar, E Ionides, C. Codeço, and M Pascual (2020). Predicting re-emergence times of dengue epidemics at low reproductive numbers: DENV1 in Rio de Janeiro, 1986–1990. Journal of the Royal Society Interface 17, 20200273.

Taylor DJ, K Dittmar, MJ Ballinger, and JA Bruenn (2011). Evolutionary maintenance of filovirus-like genes in bat genomes. BMC Evolutionary Biology 11, 1–12.

Thorson A, P Formenty, C Lofthouse, and N Broutet (2016). Systematic review of the literature on viral persistence and sexual transmission from recovered Ebola survivors: evidence and recommendations. BMJ open 6, e008859.

Towner JS, BR Amman, TK Sealy, SAR Carroll, JA Comer, A Kemp, R Swanepoel, CD Paddock, S Balinandi, ML Khristova, PBH Formenty, CG Albarino, DM Miller, ZD Reed, JT Kayiwa, JN Mills, DL Cannon, PW Greer, E Byaruhanga, EC Farnon, P Atimnedi, S Okware, E Katongole-Mbidde, R Downing, JW Tappero, SR Zaki, TG Ksiazek, S. Nichol, and PE Rollin (July 2009). Isolation of Genetically Diverse Marburg Viruses from Egyptian Fruit Bats. en. PLOS Pathogens 5. Publisher: Public Library of Science, e1000536. ISSN: 1553-7374. 10.1371/journal.ppat.1000536.

Zhao S, Y Lou, AP Chiu, and D He (2018). Modelling the skip-and-resurgence of Japanese encephalitis epidemics in Hong Kong. Journal of theoretical biology 454, 1–10.

